## Supplementary information for "Large protein databases reveal structural complementarity and functional locality"

Paweł Szczerbiak, Lukasz Szydlowski, Witold Wydmański,
P. Douglas Renfrew, Julia Koehler Leman, Tomasz Kosciolek*

**Structural clustering** 1
**First stage: MIP database** 2
**The largest cluster of MIP novel folds** 4
**First stage: highquality_clust30 dataset** 6
**Second stage: final dataset** 9
**Structure space** 14
**PaCMAP grid search** 14
**Visualizations** 18
**Plots for unnormalized Geometricus representations** 19
**Other databases** 21
**Functional annotations** 27
**deepFRI v1.1** 27
**Validation using E. coli proteome** 27
**Visualizations** 29
**Plots for unnormalized Geometricus representations** 30
**Top COG categories** 32
**Cluster heterogeneity** 33
**Taxonomy analysis** 40
**References** 42

### Structural clustering

Similarly to reference (1) we used Foldseek to remove structural redundancy and find cluster representatives. In all cases, we used cov-mode = 0 (coverage of query and target). By cluster we understand a group of similar structures of cardinality at least 2 (the rest are singletons). We utilized a two-stage procedure (see Fig. 1A in the main text):

- First stage: cluster each dataset independently i.e. AFDB50 (already done in reference (1)), highquality_clust30 (high quality predictions from ESMAtlas with pTM and pLDDT > 0.7 clustered at 30% sequence similarity level) [https://github.com/facebookresearch/esm/blob/main/scripts/atlas/README.md], and MIP (2) (mostly single-domain structures already filtered at 30% sequence identity level). Foldseek parameters (e-value and coverage) have been tuned separately for each dataset (see subsequent paragraphs).

- Second stage: gather cluster representatives and MIP singletons (we exclude AFDB and ESMAtlas singletons from the reasons described in the main text – see Methods section), and cluster this set with Foldseek.

### First stage: MIP database

To find optimal Foldseek parameters for the MIP database, which comprises short (between 40 and 200 residues) mostly single-domain proteins, we chose e-values = [0.1, 0.01, 0.001, 0.0001] and structural alignment overlap c = [0, 0.9]. The second value for the c parameter was chosen to align with the procedure in reference (1). For similar reasons, we did not consider more extreme e-values.

Non-zero coverage parameter c results in more clusters (and singletons) since we require that two given structures must be not only similar but also of comparable size – see Fig. S1. It is also clear from the plot that the number of singletons scales almost linearly with e-value. For the number of clusters it is more complicated but we can notice that e-value = 0.001 provides the biggest number of clusters for c = 0; for c = 0.9 we can observe a small decrease with e-value with a sharp drop at 0.0001. As a consequence, mean cluster size also decreases with e-value and c parameter – see Fig. S2. The same concerns the maximum cluster size for c = 0.9; interestingly, for c = 0 we can notice an increase in e-value with a large drop, again at 0.0001. Those effects are probably heavily dataset-dependent but, in general, e-values between 0.1 and 0.001 should provide a reasonable tradeoff between the number of clusters and their size. In Fig. S3 we show that a non-zero c parameter is critical if we need to ensure cluster consistency. For c = 0 representative structures might be short (two times smaller in this case) and not grasp the full diversity of the clusters they represent. Interestingly, for c = 0.9 the distribution of the longest structure size to representative structure size almost does not depend on e-value.

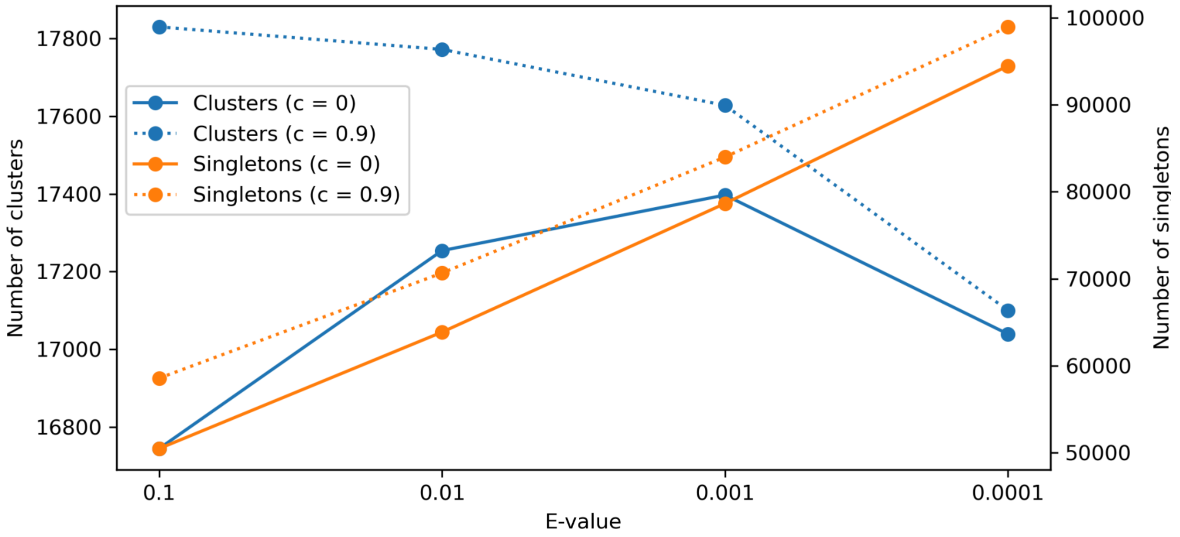

**Fig. S1:** Number of clusters (left y-axis) and singletons (right y-axis) as a function of e-value stratified by coverage parameter, c.

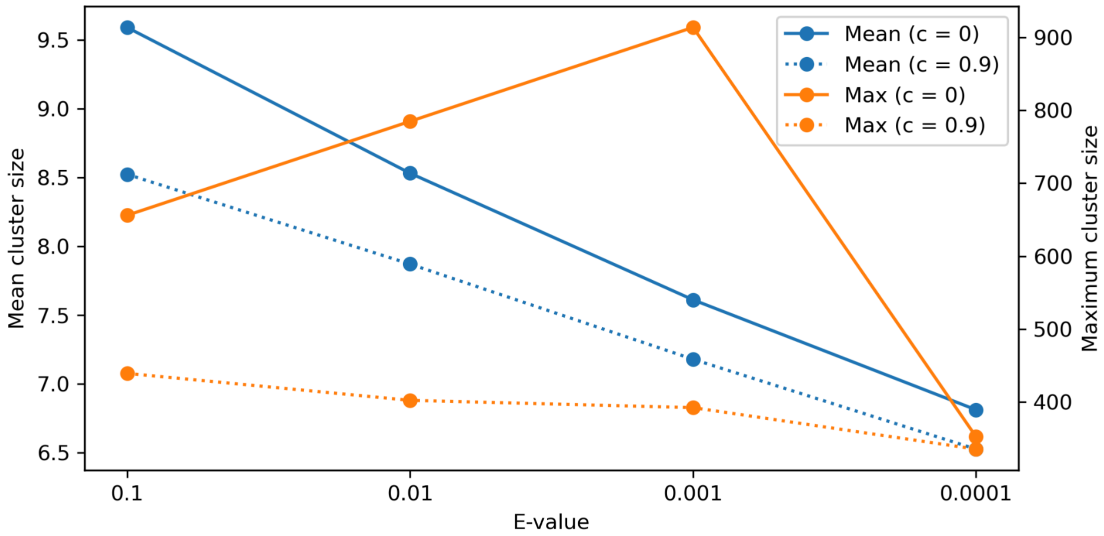

**Fig. S2:** Mean cluster size (left y-axis) and maximum cluster size (right y-axis) as a function of e-value stratified by coverage parameter, c. Note that the standard deviation is not meaningful here because the distribution of cluster sizes does not follow a normal distribution.

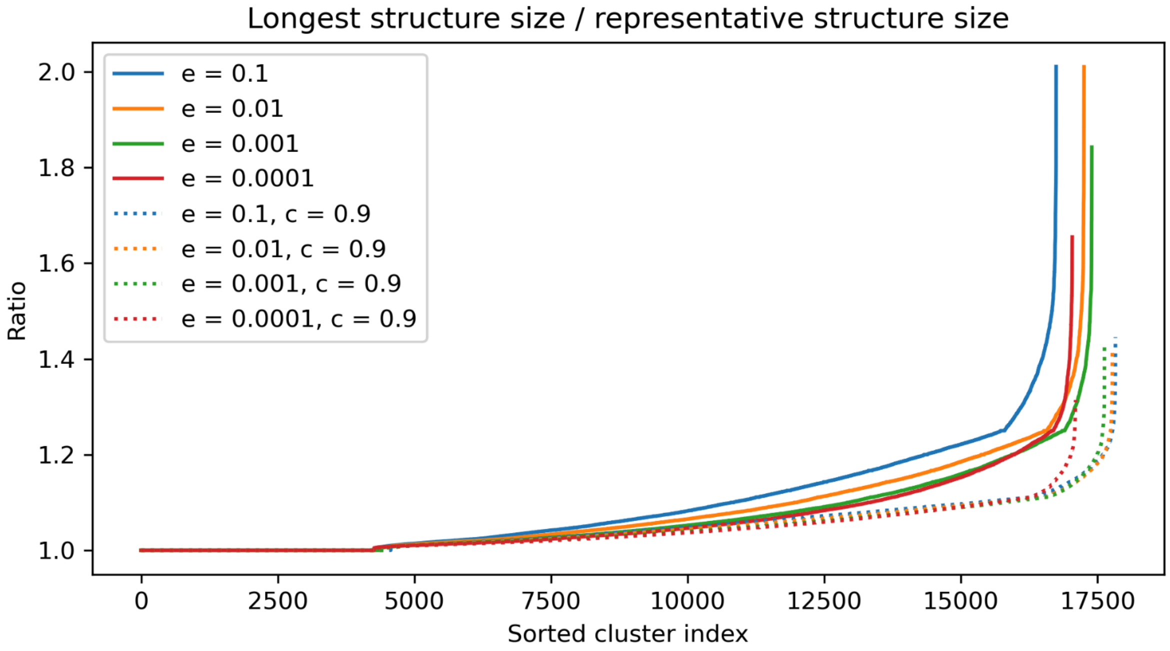

**Fig. S3:** Ratio (y-axis) between the longest and representative structure sizes for a given cluster (x-axis). Cluster indices in the x-axis are sorted based on the value on the y-axis. Each line corresponds to a different e-value, e, and coverage parameter, c.

To estimate structural similarity across clusters we superimposed all structures within each cluster with US-align – see Fig. S4 and Fig. S5. According to expectations, the TM-score increases with e-value, however, the differences are not large (at least for e-value ≥ 0.01). Interestingly, coverage, c, does not influence structural similarity so much in this case. Still, we can observe a larger discrepancy between minimum, mean, and maximum TM-score distribution for c = 0 as compared to c = 0.9.

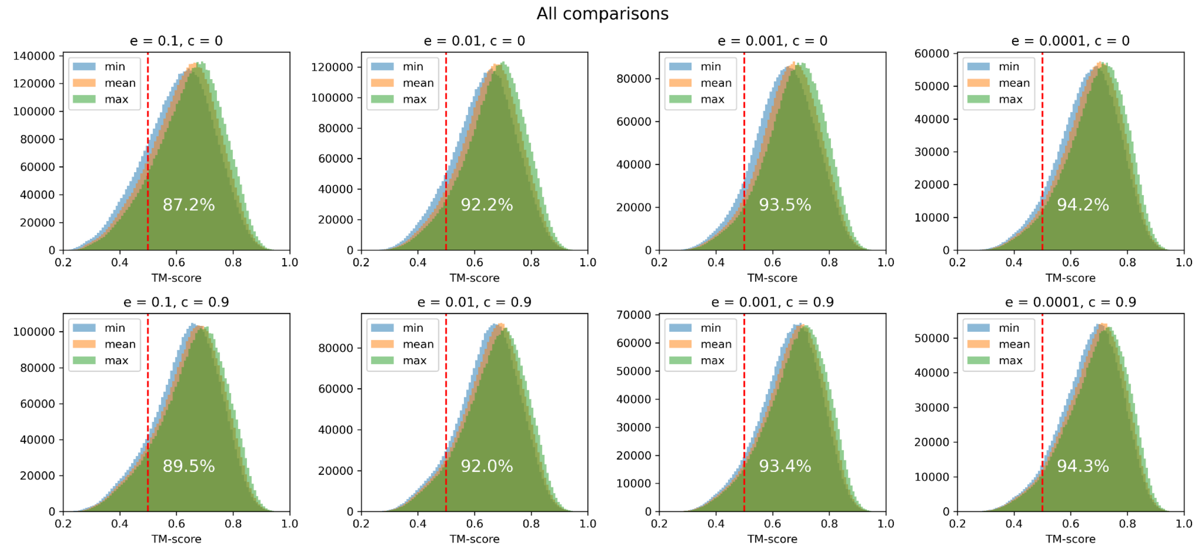

**Fig. S4:** Distribution of minimum, mean, and maximum TM-score between each two structures (excluding identities) within a given cluster for all clusters. Each panel corresponds to a different e-value, e, and coverage parameter, c, combination. Numbers in white denote percentages of points with mean TM-score ≥ 0.5 (red vertical line).

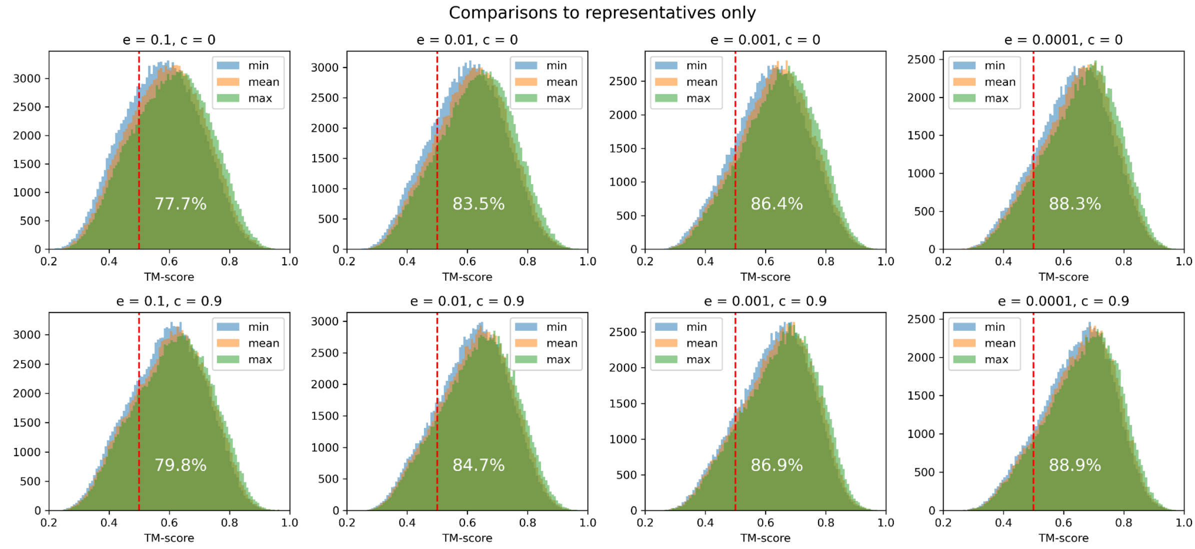

**Fig. S5:** Distribution of minimum, mean, and maximum TM-score between each structure and representative structure (excluding identities) within a given cluster for all clusters. Each panel corresponds to a different e-value, e, and coverage parameter, c, combination. Numbers in white denote percentages of points with mean TM-score ≥ 0.5 (red vertical line).

#### The largest cluster of MIP novel folds

As a final verification of foldseek clustering, we checked whether we could identify the largest cluster of MIP novel folds (Supplementary Figure 53 in reference (2)). Surprisingly, for each combination of foldseek parameters that we tested (most importantly, all e-values), the cluster comprised 105 structures including all 87 found in the aforementioned MIP cluster – see Fig. S6. What is even more interesting, the additional 18 structures demonstrate very high TM-score within themselves and against the 87 ones (Fig. S7). **It proves that Foldseek is indeed a reliable protein structure clustering method.** The reason why those additional structures had not been taken into account when constructing the MIP novel folds is the prefiltering step. Namely, we required the DMFold and Rosetta models to both have max TM-score against PDB90 ≥ 0.5 which was not fulfilled (sometimes at a marginal level e.g. 0.01). This indicates that using rigid thresholds on similarity metrics for identifying potential novel folds may not be the most advantageous approach.

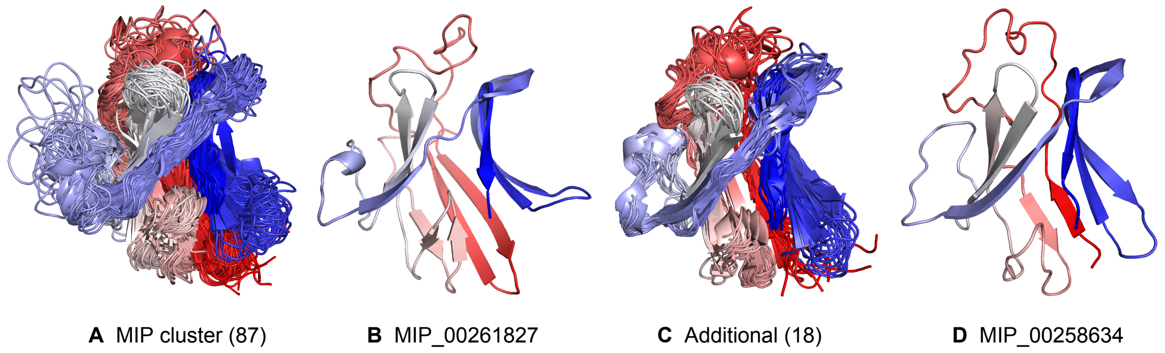

**Fig. S6:** Foldseek cluster that overlaps with the largest cluster of MIP novel folds. (**A**)**:** Structures that have been found also in the MIP cluster. (**B**)**:** Example structure in **A**. (**C**)**:** Additional structures found in the Foldseek cluster. (**D**)**:** Example structure in **A**.

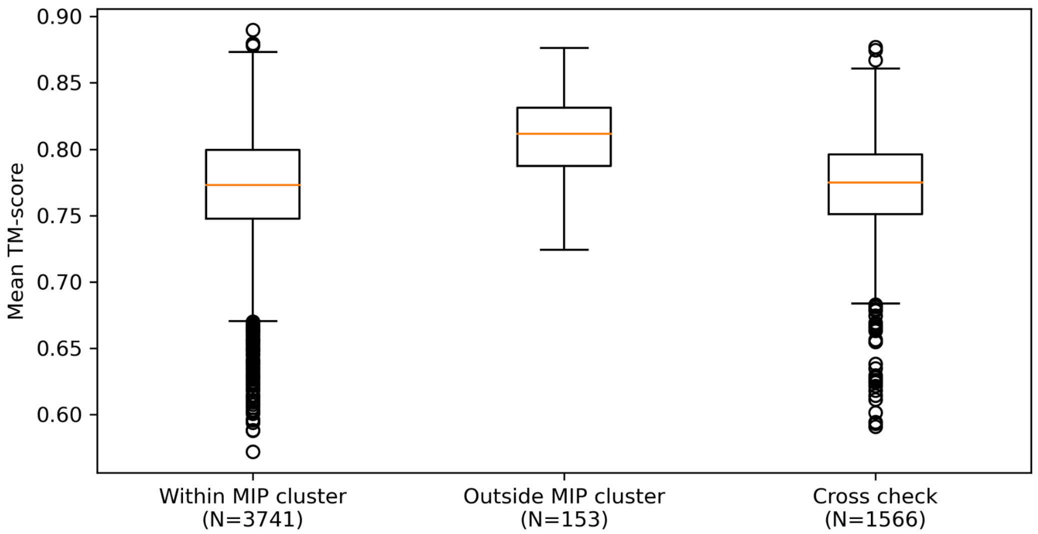

**Fig. S7:** Mean TM-score between all superimposed structures within a given group i.e. (from left to right) superposition of structures present in the largest cluster of MIP novel folds (Fig. S6A), superposition of additional structures not present in the previous group (Fig. S6C), crossed superposition between first and second group. Number of data points for each box is shown below the x-axis.

In summary, the optimal set of analyzed hyperparameters that provides high consistency and similarity within clusters and a satisfactory number of clusters/singletons for the MIP database is **e-value = 0.001** and **c = 0.9**. The key observation is, however, that many features (including identification of compact clusters) do not depend so strongly on e-value.

#### First stage: highquality_clust30 dataset

Similarly, we utilized the same procedure to cluster a high-quality subset of ESMAtlas, i.e. highquality_clust30. Results are presented in plots Fig. S8–Fig. S10. We might notice many similarities to the MIP database clustering. Interestingly, the mean cluster size (which can be treated as a measure of structural redundancy) is a few times larger as compared to the MIP.

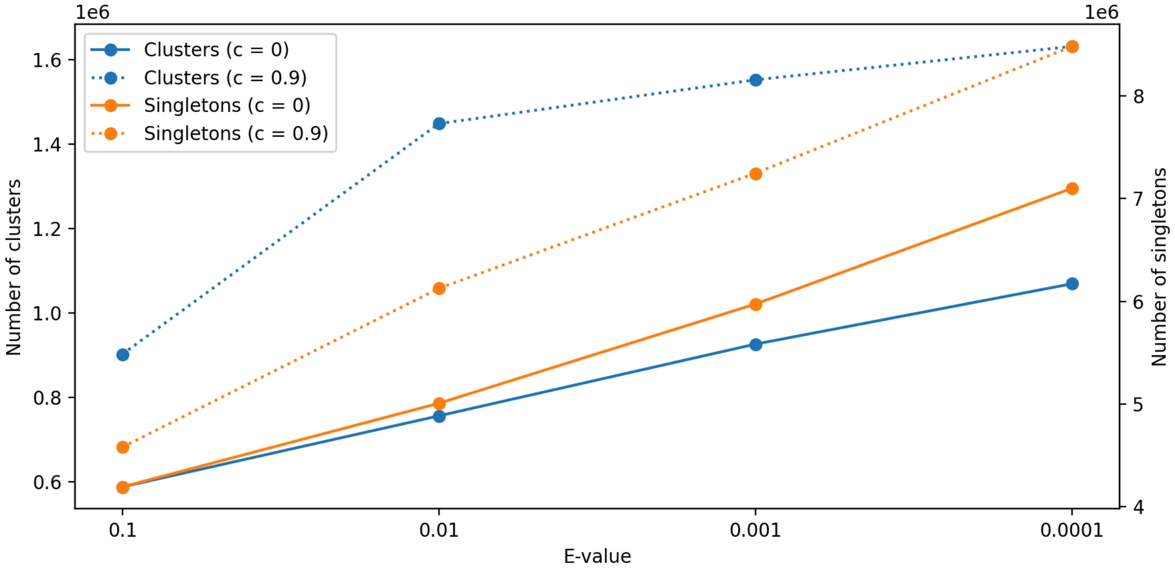

**Fig. S8:** Number of clusters (left y-axis) and singletons (right y-axis) as a function of e-value stratified by coverage parameter, c.

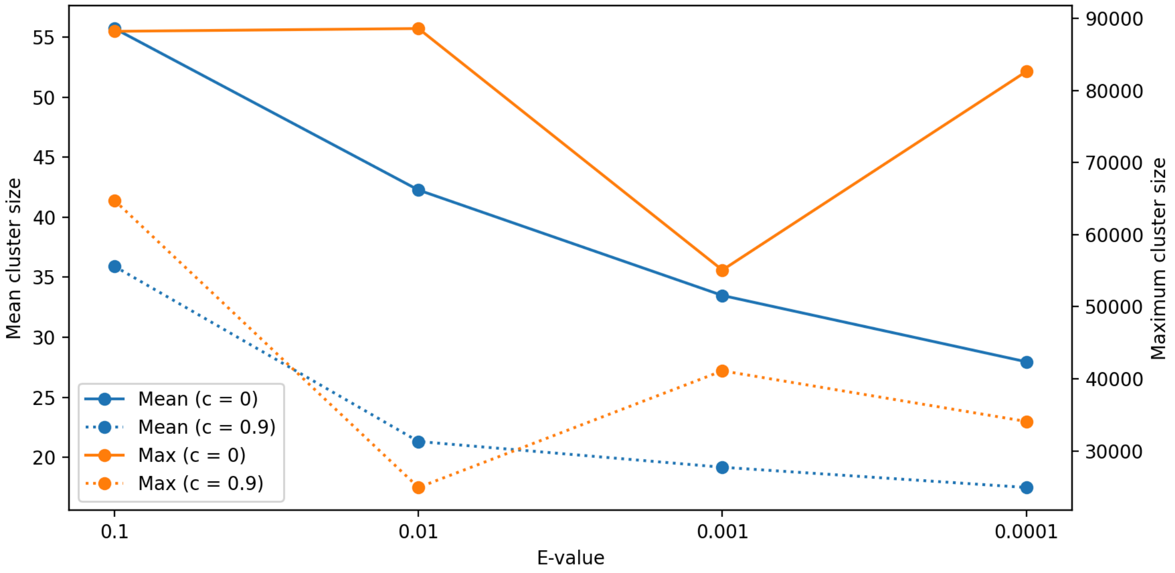

**Fig. S9:** Mean cluster size (left y-axis) and maximum cluster size (right y-axis) as a function of e-value stratified by coverage parameter, c. Note that the standard deviation is not meaningful here because the distribution of cluster sizes does not follow a normal distribution.

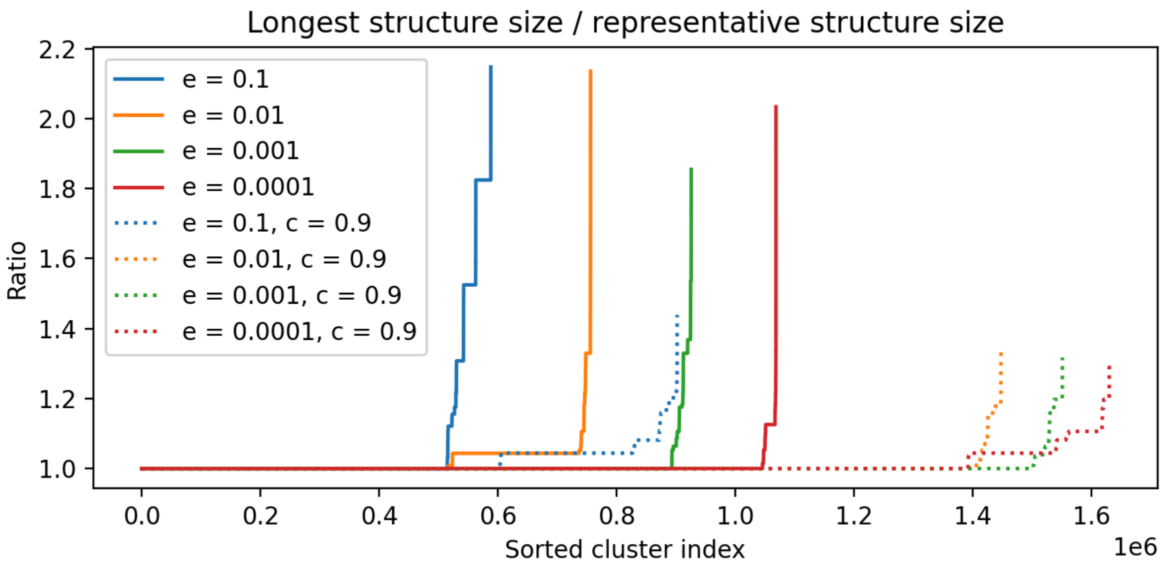

**Fig. S10:** Ratio (y-axis) between the longest and representative structure sizes for a given cluster (x-axis). Cluster indices in the x-axis are sorted based on the value on the y-axis. Each line corresponds to a different e-value, e, and coverage parameter, c.

For further investigation, we chose three combinations i.e. e-value = 0.01, 0.001, 0.0001, and c = 0.9. When it comes to the overall TM-score distribution (see Fig. S11), all e-values provide satisfactory outcomes. Surprisingly, the TM-score between the representative and all the other structures for the top 12 largest clusters favors e-value = 0.01 (see Fig. S12). In general, however, the number of poor (median TM-score between representative structure and all the other structures per cluster between 0.4 and 0.5) and very poor clusters (median TM-score smaller than 0.4) decreases with increasing e-value – see Fig. S13 and Table S1.

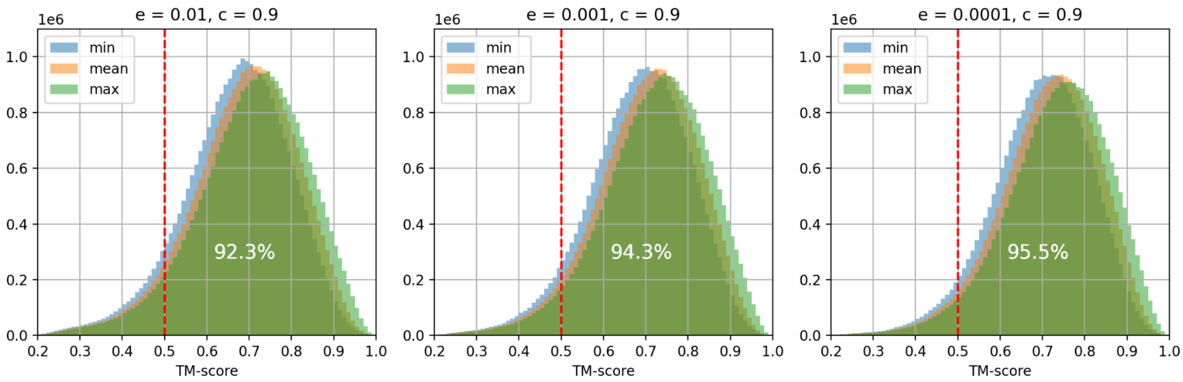

**Fig. S11:** Distribution of minimum, mean, and maximum TM-score between each structure and representative structure (excluding identities) within a given cluster for all clusters. Each panel corresponds to a different e-value, e, and coverage parameter, c, combination. Numbers in white denote percentages of points with mean TM-score ≥ 0.5 (red vertical line).

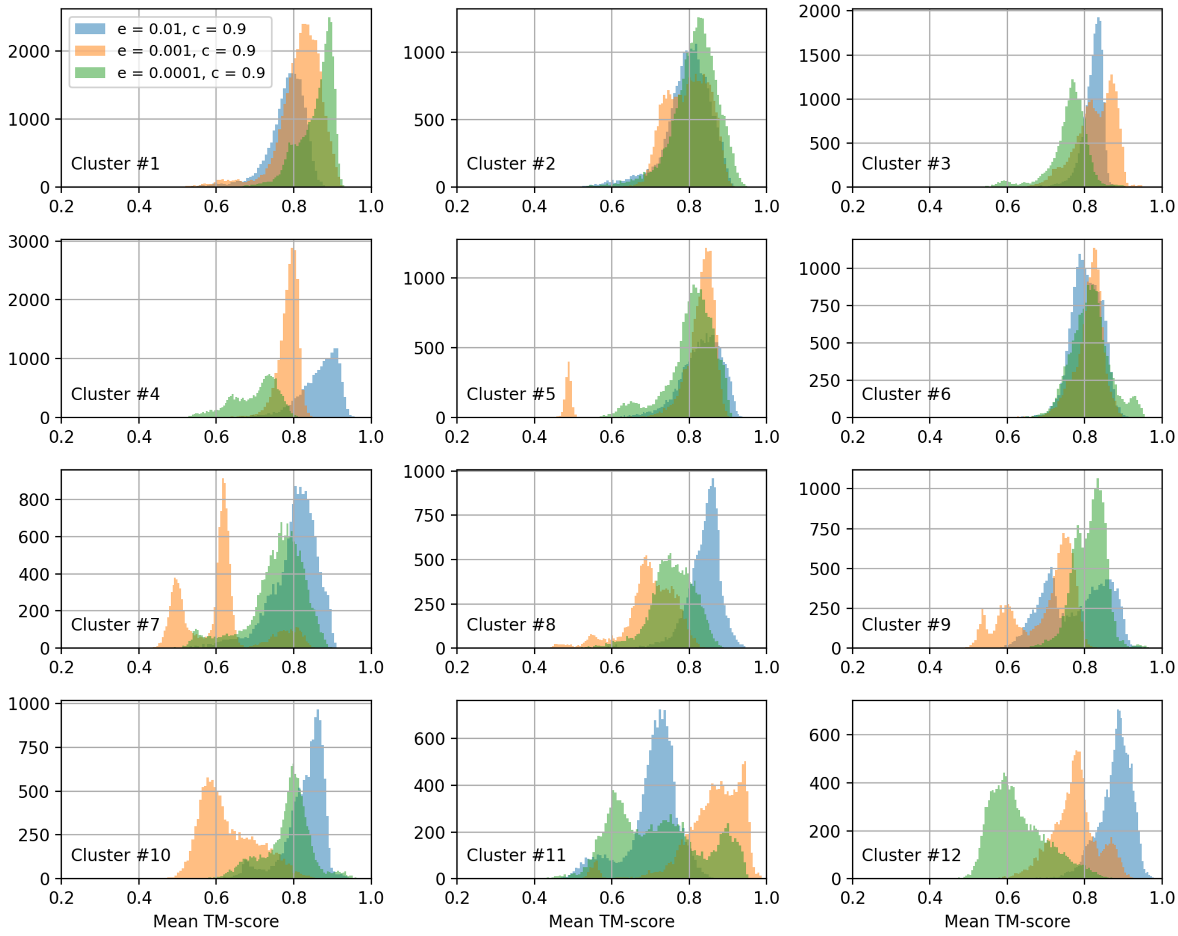

**Fig. S12:** Distribution of mean TM-score between each structure and representative structure (excluding identities) within a given cluster for the top 12 largest clusters. Each color corresponds to a different e-value, e, and coverage parameter, c, combination (see top left legend).

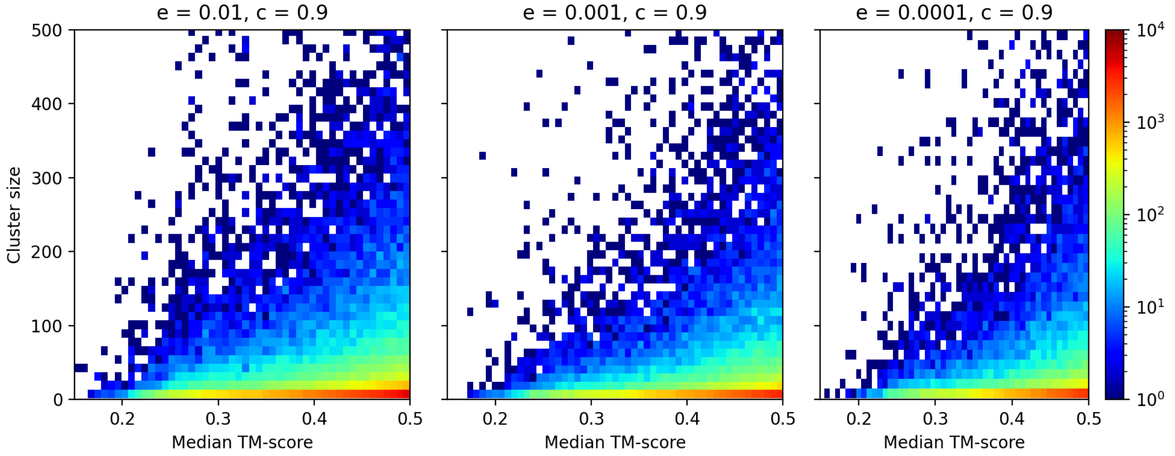

**Fig. S13:** Heatmap between median TM-score (chosen between mean TM-scores of cluster representative to all the other structures in a given cluster) and cluster size. Each panel corresponds to a different e-value, e, and coverage parameter, c, combination. In contrast to Fig. S12 we focused on small and medium clusters (between 2 and 500 elements) with poor quality (TM-score < 0.5).

**Table S1:** Number of very poor large clusters (size > 500, median TM-score between representative structure and all the other structures per cluster < 0.4). Coverage parameter c = 0.9.

| **E-value** | 0.01 | 0.001 | 0.0001 |
|---|---|---|---|
| **Number of clusters** | 71 | 44 | 29 |

Taking all the above into consideration (especially the best cluster quality among all combinations and the largest number of cluster representatives), we chose **e-value = 0.0001 and c = 0.9** as the optimal clustering parameters.

### Second stage: final dataset

Finally, to remove structural redundancy among different, already structurally clustered, datasets, we used Foldseek yet again but this time for coverage parameter c = 0.7, 0.8, and 0.9 (following the work of Barrio-Hernandez et al. or standards set by UniRef (1, 3)). Overall (except c = 0.9, e-value = 0.0001), the number of clusters and singletons (mean cluster size) increases (decreases) with e-value and coverage parameter, c – Fig. S14 and Fig. S15. Interestingly, the number of singletons for c = 0.9 is ~2 times larger as compared to c = 0.7 and 0.8.

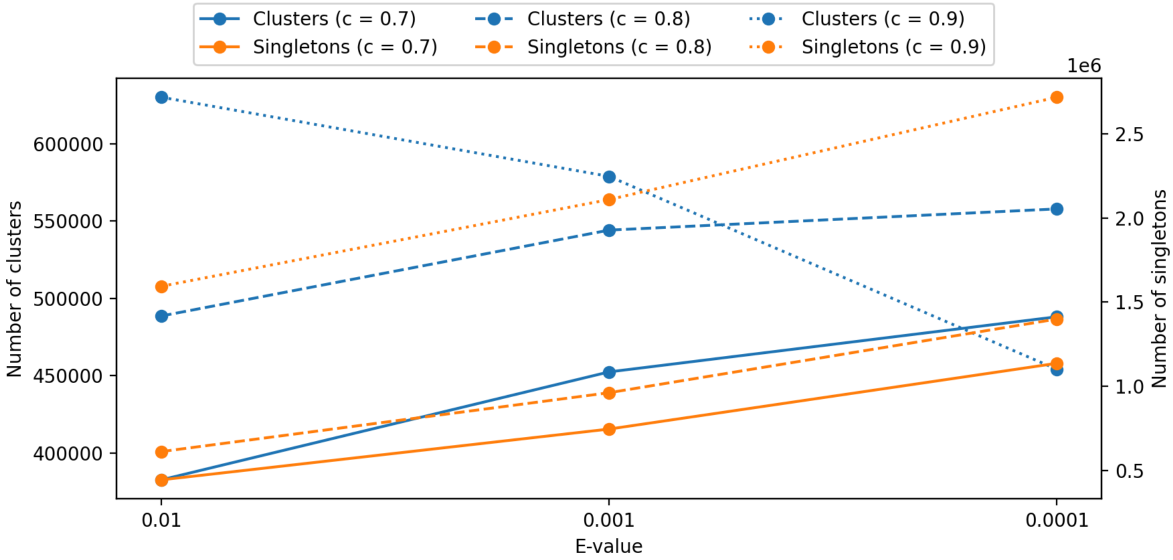

**Fig. S14:** Number of clusters (left y-axis) and singletons (right y-axis) as a function of e-value stratified by coverage parameter (c).

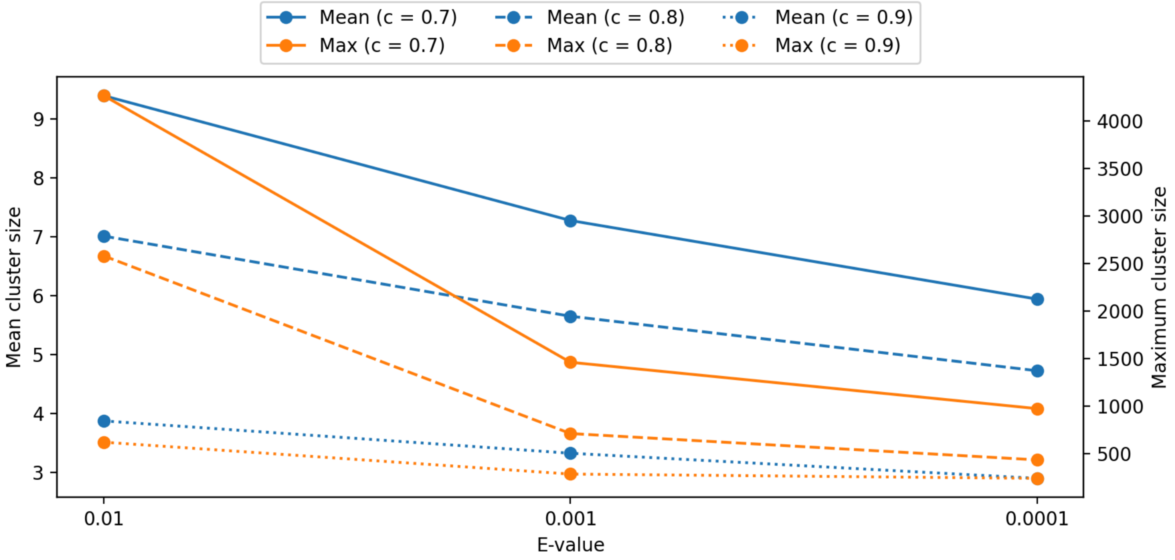

**Fig. S15:** Mean cluster size (left y-axis) and maximum cluster size (right y-axis) as a function of e-value stratified by coverage parameter (c). Note that the standard deviation is not meaningful here because the distribution of cluster sizes does not follow a normal distribution.

The TM-score distribution between the representative and all the other structures for the top 10 largest clusters shows that c = 0.7 is too permissive (see Fig. S16).

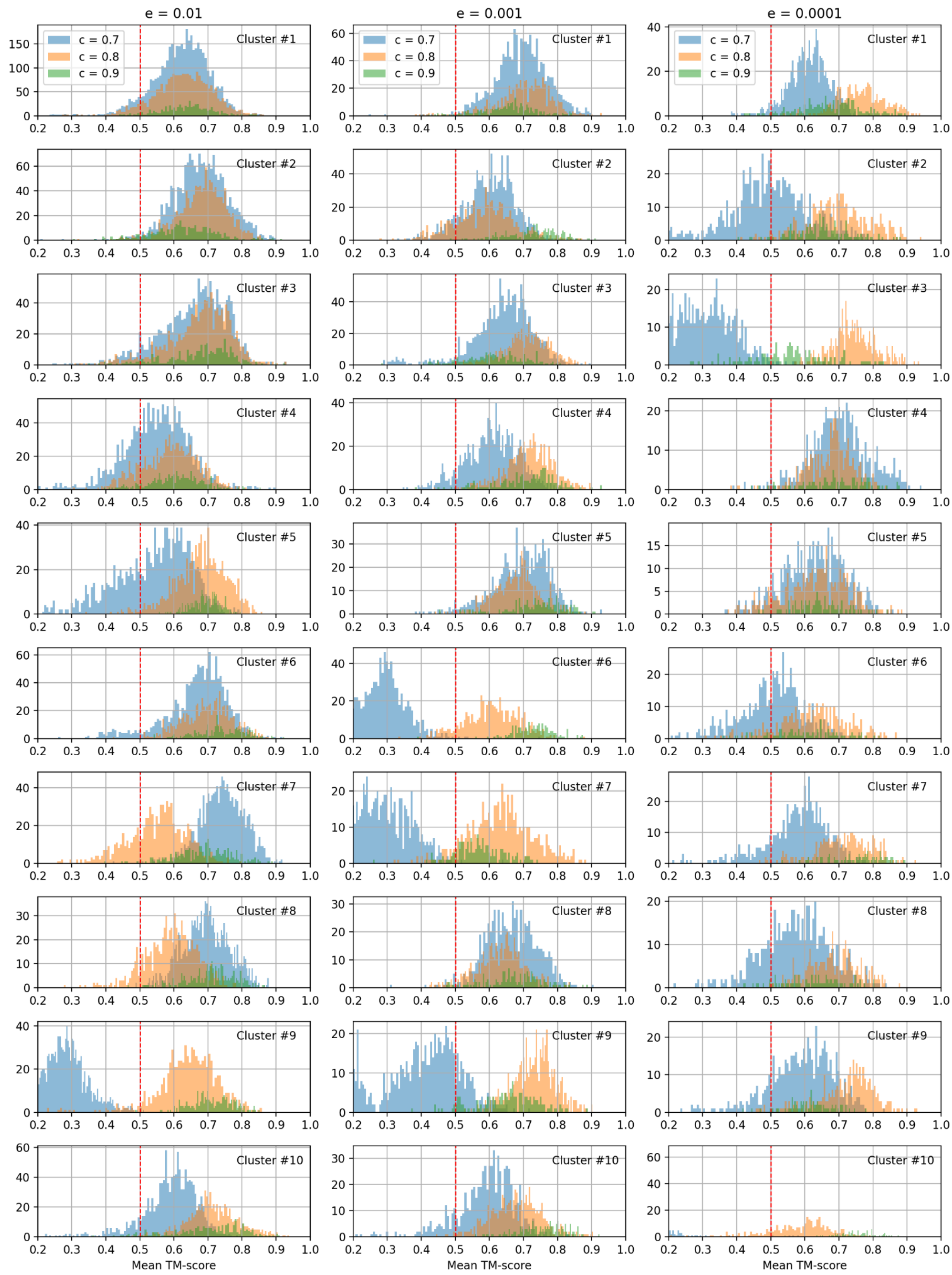

**Fig. S16:** Distribution of mean TM-score between each structure and representative structure (excluding identities) within a given cluster for the top 10 largest clusters. Each color corresponds to a different coverage parameter, c, whereas each column to a different e-value, e.

In summary, the optimal combination, providing a reasonable number of cluster representatives including singletons (reduction from 4,035,121 to 1,505,141) and high-quality clusters, is **e-value = 0.001 and c = 0.8**. Smaller/larger e-values would also give satisfactory results (contrary to the coverage parameter).

Table S2 summarizes the number of structures in each clustered dataset (including final dataset). Note that we consider here all AFDB structures (including those with low pLDDT). Restricting to high-quality (pLDDT > 70) left us with ~62% light clusters and only 48% of dark clusters. Fig. S17 shows structure length distribution including only high-quality predictions. Fig. S18 presents distribution of mean pLDDT for AFDB structures from the full database (random samples) and clustered dataset. Clearly, we can notice a huge discrepancy between the two, caused by clustering procedures (many high-quality models are similar and have been removed due to redundancy).

**Table S2:** Optimal Foldseek parameters, number of input structures, and the resulting number of clusters/singletons for each dataset (first stage clustering, first four rows). In yellow/red we indicate representative structures that have been used in/excluded from the second stage clustering (last row). In green we present final numbers of structures that are analyzed in this work. Number of input structures refers to non-redundant structures on a sequence level. Number of output structures refers to all structures gathered in the clusters (and singletons, but only for MIP and the final database). For the final database we show numbers of all output clusters, singletons, and structures as well as only the high-quality ones (i.e., with mean pLDDT > 70 for AFDB structures; other databases are already considered high-quality).

| | | Foldseek e-value | Foldseek coverage | Input structures | Clusters | Singletons | Output structures |
|---|---|---|---|---|---|---|---|
| **AFDB light*** | | 0.01 | 0.9 | 52,327,413 | 1,591,199 | 13,012,338 | 30,045,210 |
| **AFDB dark*** | | | | | 711,700 | | |
| **MIP** | | 0.001 | 0.9 | 211,069 | 17,628 | 83,986 | 211,069 |
| **ESMAtlas** | | 0.0001 | 0.9 | 36,983,470 | 1,630,608 | 8,482,776 | 28,500,694 |
| **Final database** | **all** | 0.001 | 0.8 | 4,035,121 | 544,123 | 961,018 | 4,035,121 |
| | **hq** | | | | 422,003 | 648,398 | 3,060,808 |

*Clustering performed in reference (1).

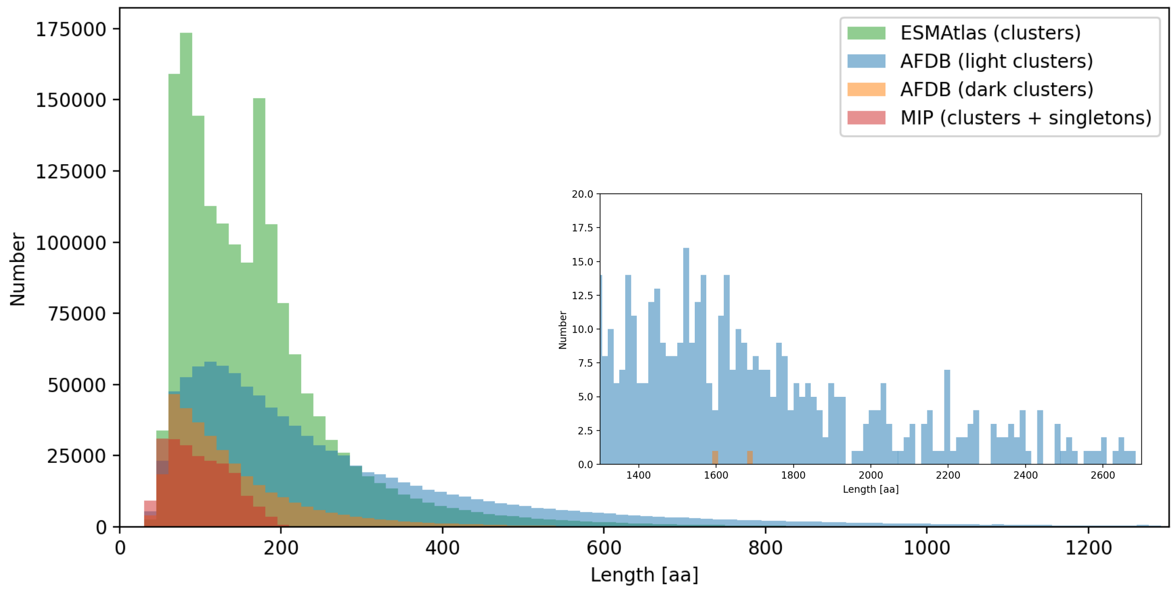

**Fig. S17:** Structure length distribution in the analyzed datasets (only high-quality predictions have been considered – see last row in Table S2). Inset shows long proteins (not visible in the main panel).

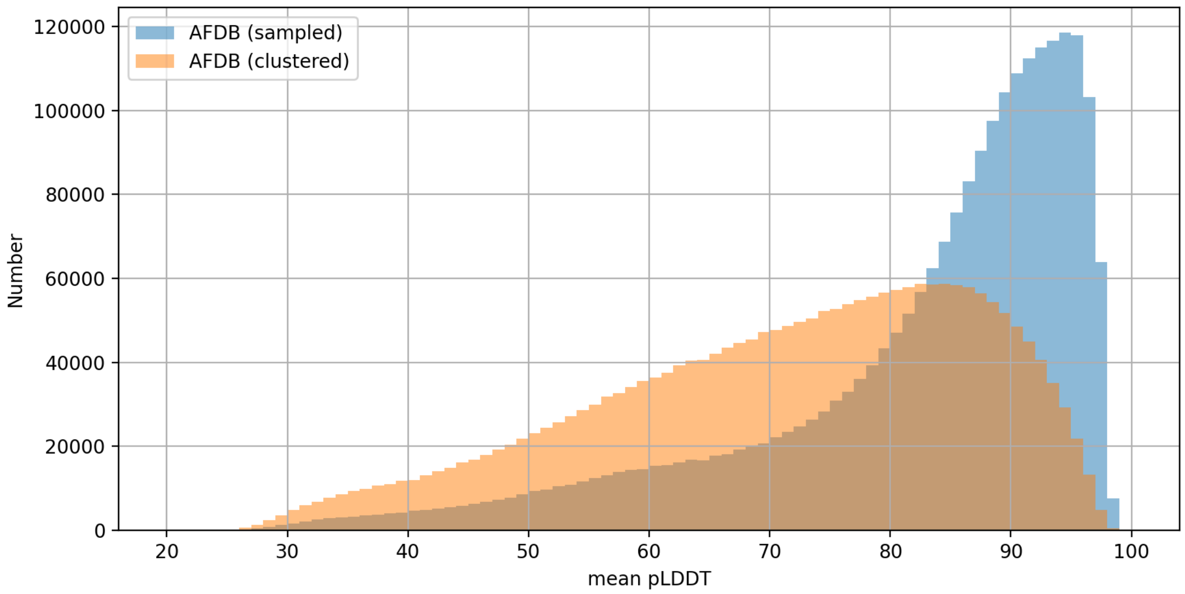

**Fig. S18:** Histogram of mean pLDDT for AFDB structures: randomly sampled from the entire database and coming from the clustered dataset. Both distributions have equal sample sizes for accurate comparison.

### Structure space

Shape-mer representations generated with Geometricus for all representative structures in the final clustered database have been reduced into two dimensional space using PaCMAP (see Methods in the manuscript). We considered two versions of embedding vectors:

- unnormalized: raw shape-mer vector (summing up to the number of residues of a query protein)
- normalized: raw shape-mer vector divided by the sum of its elements (summing up to 1)

Of course, the second type is less biased by the structure length distribution so we chose it as a default.

#### PaCMAP grid search

In order to find optimal PaCMAP parameters we performed a grid search, changing:

- `n_neighbors` between 2 to 20 every 1
- `MN_ratio` between 0.1 to 5.1 every 0.2
- `FP_ratio` between 0.1 to 5.1 every 0.2

We wanted to ensure the best separation between clusters from Fig. S16 but 9 out of 10 cluster representatives are alpha-like and one is alpha/beta-like. Therefore, we decided to add 5 additional clusters that are beta- and alpha/beta-like (see Fig. S19). This yielded 8,180 structures, for which we used the agreement TM-score as a measure of similarity.

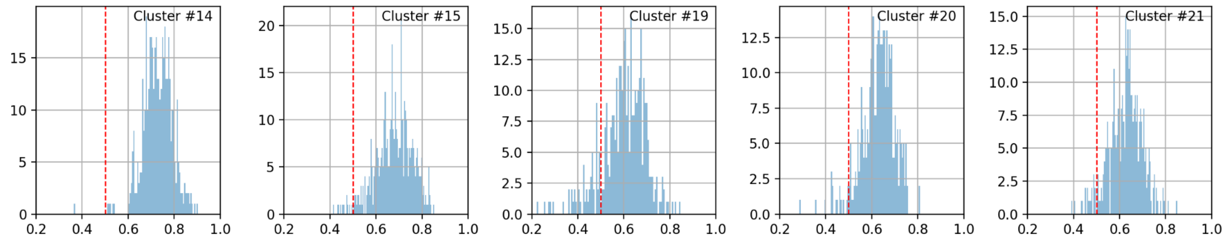

**Fig. S19:** The same as in Fig. S16 but for e-value = 0.001, c = 0.8 and for manually chosen beta- and alpha/beta-like clusters.

We wanted to preserve both global (between clusters) and local (within a cluster) separation. For each PacMAP reduction we computed correlation (Pearson and Spearman coefficients) between Euclidean distance for each pair of points (structures) and their mean TM-score, taking two inputs:

- representative structures – see Fig. S20
- all structures from each cluster (we take mean correlation coefficient as a final outcome) – see Fig. S21 and Fig. S22

Our maximization function is simply a product of the two coefficients defined above, separately for Pearson and Spearman – note that both are negative so their product is positive. Fig. S23 gathers the results.

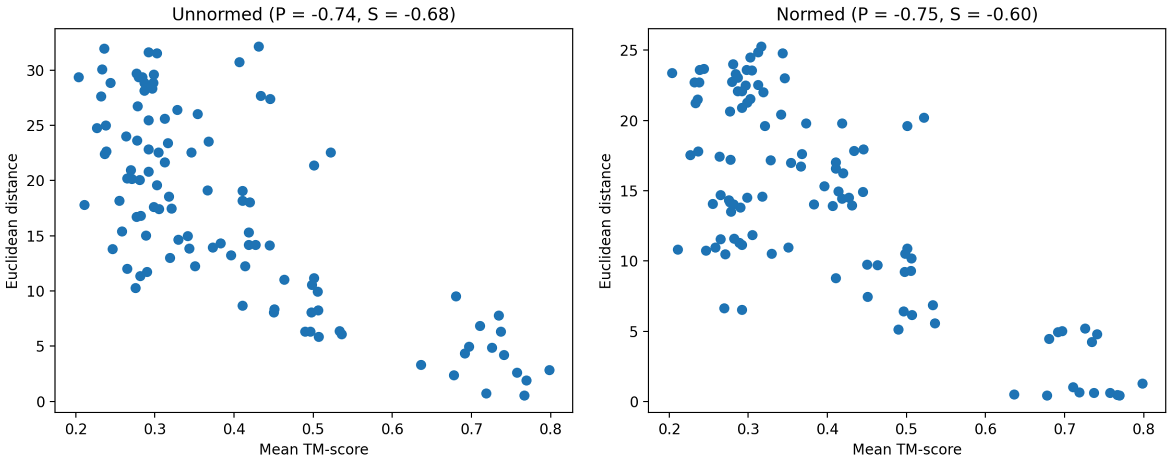

**Fig. S20:** Mean TM-score versus Euclidean distance for representative structures. Left/right: unnormalized/normalized shape-mer representation. In the title P and S denote Pearson and Spearman correlation coefficients respectively.

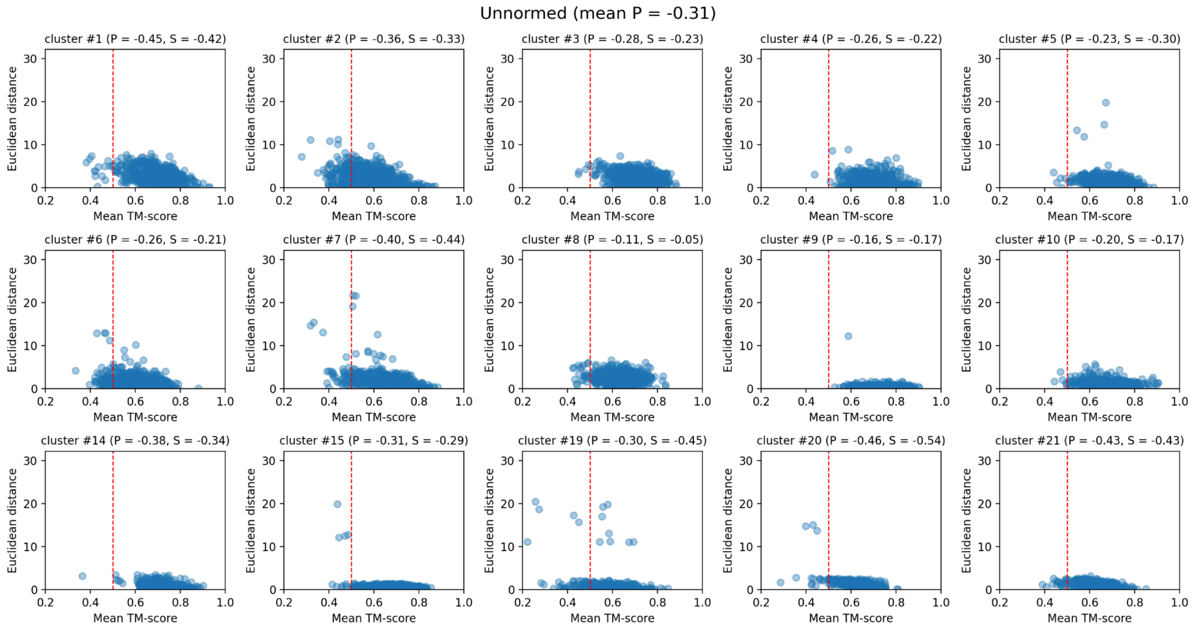

**Fig. S21:** Mean TM-score versus Euclidean distance for all structures within a given cluster for unnormalized shape-mer representation. In the titles P and S denote Pearson and Spearman correlation coefficients respectively.

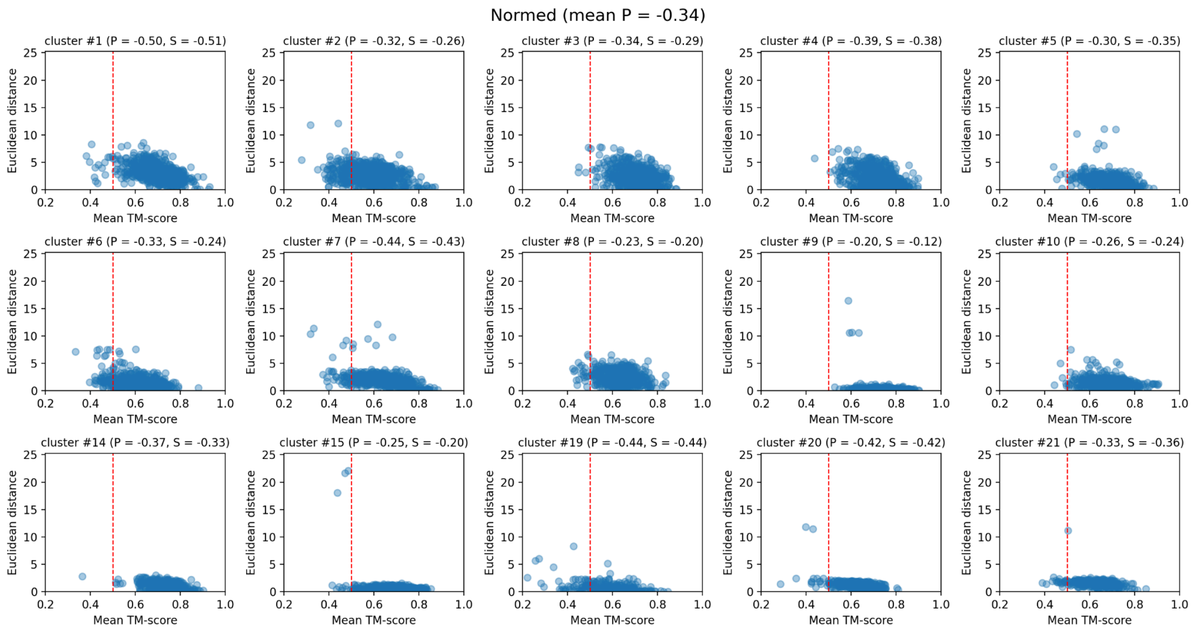

**Fig. S22:** Mean TM-score versus Euclidean distance for all structures within a given cluster for normalized shape-mer representation. In the titles P and S denote Pearson and Spearman correlation coefficients respectively.

There are a few comment worth noting:

- In general, larger Spearman and Pearson products are obtained for normalized Geometricus representations (panel A in Fig. S23).
- We can observe certain relations between `MN_ratio` and `FP_ratio`, and default parameters may not always be optimal (panel B in Fig. S23).
- Default value of `n_neighbors` seems to be quite robust and in our case any value equal or larger than 10 gives good performance (panel C in Fig. S23).

In summary, the following sets of optimal PaCMAP parameters (`n_neighbors`, `MN_ratio`, `FP_ratio`) has been chosen:

- unnormalized: (13, 1.9, 1.5)
- normalized: (10, 1.3, 0.9)

Fig. S24 and Fig. S25 illustrate the PaCMAP reduction in action, showing a clear separation between alpha and beta clusters. Notably, within the alpha clusters, which overrepresent the top largest clusters, there is a distinct arrangement reflecting finer details of the folds. Obviously, for the unnormalized Geometricus vectors, proteins are distributed according to structure length, a pattern not observed with the normalized inputs.

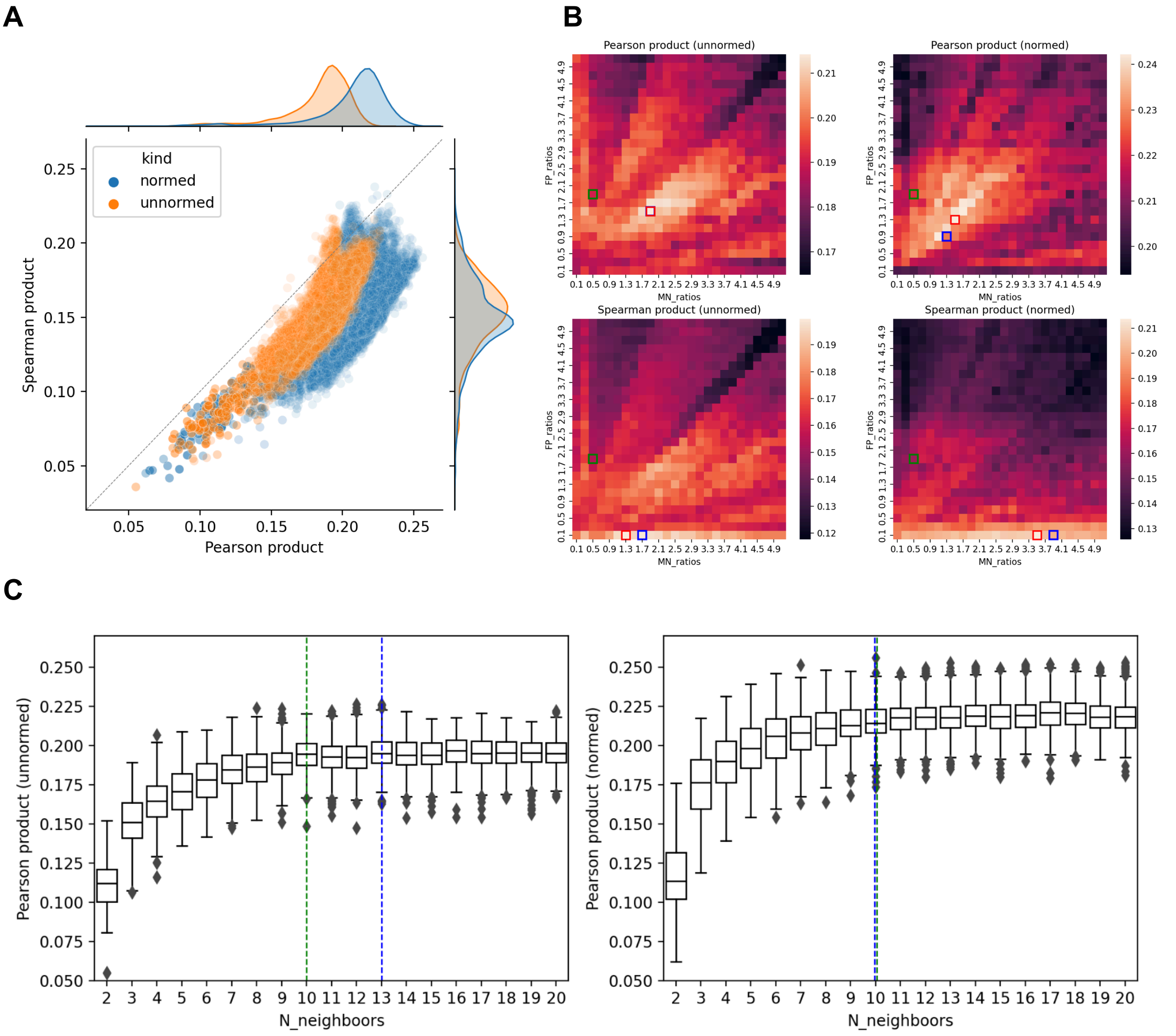

**Fig. S23: Summary of PaCMAP grid search. (A)**: Scatter plot of Pearson and Spearman products (see the text for definitions) for two versions of shape-mer representations (normalized and unnormalized). **(B)**: Median values (for `n_neighbors`) of Pearson and Spearman products for each tested combination of `MN_ratio` and `FP_ratio` parameters. Green/blue/red rectangles on each panel represent default/optimal for specific `n_neighbors` (the best in this case)/optimal for all `n_neighbors` (the most robust) combinations respectively. **(C)**: Dependance between Pearson and Spearman products and `n_neighbors` parameter (each box has been generated using outcomes for all `MN_ratio` and `FP_ration` parameters). Green/blue dashed lines represent default/optimal for specific `MN_ratio` and `FP_ration` (the best in this case) values respectively.

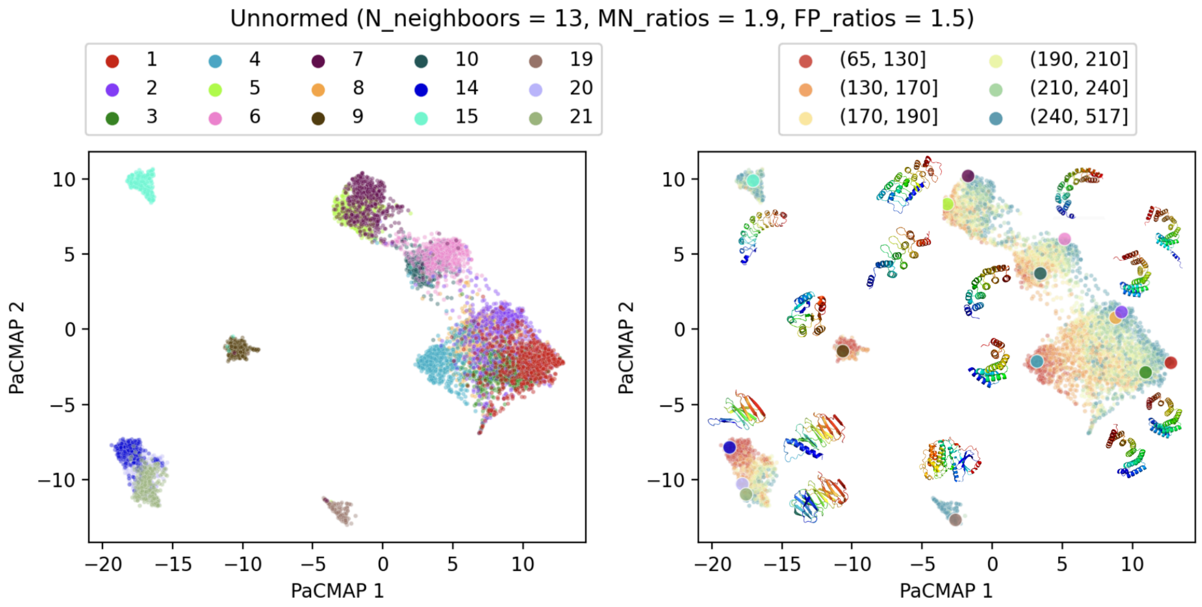

**Fig. S24:** Visualization of the protein structure space using PaCMAP (optimal parameters have been used) and unnormalized Geometricus representations for the top 15 largest clusters (see Fig. S16 and Fig. S19). Left: cluster index. Right: structure length as number of residues (see legend) and PyMOL depictions of representative structure in each cluster.

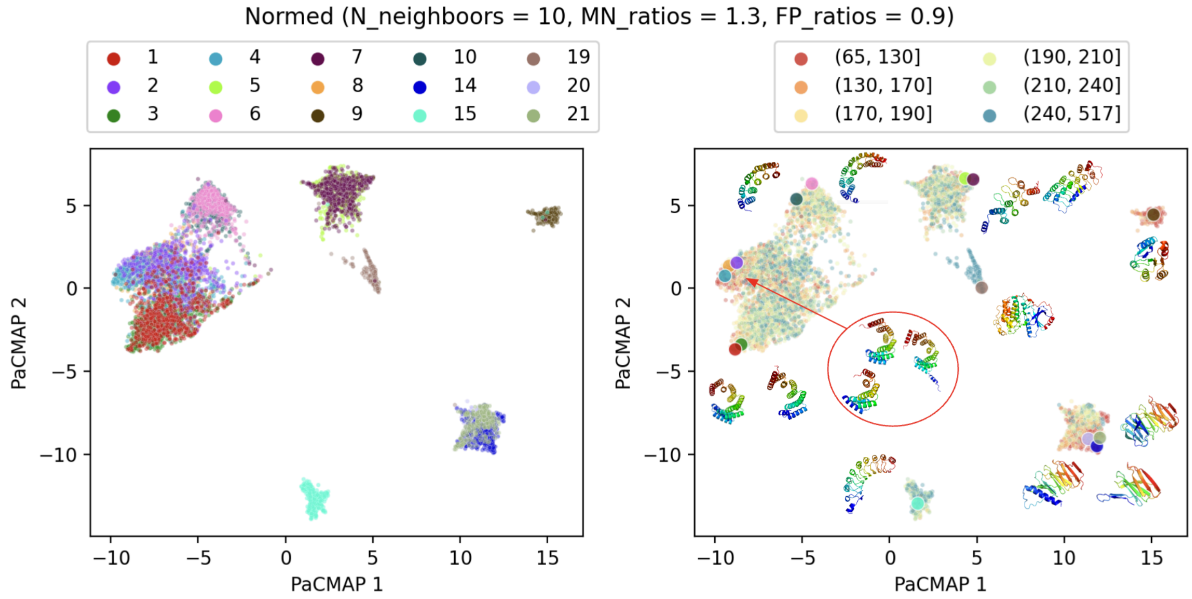

**Fig. S25:** As in Fig. S24 but for normalized Geometricus representations.

### Visualizations

Here, we provide additional panels (supplementary to the main text) for visualizing 2D protein structure space using PaCMAP, for both normalized and unnormalized Geometricus representations.

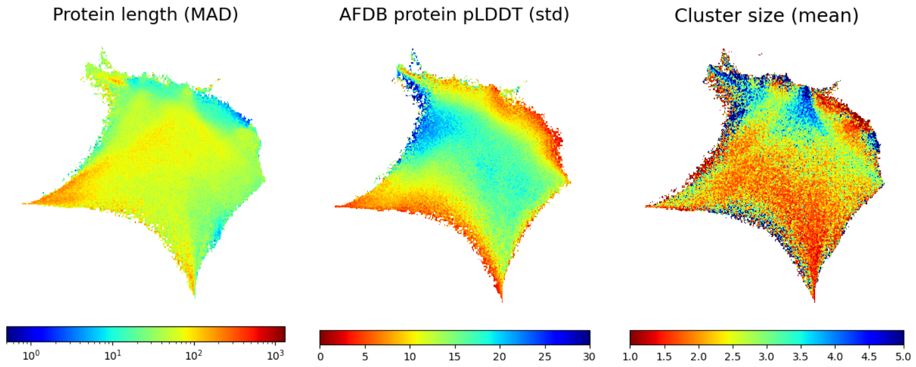

**Fig. S26:** Dispersion (MAD: median absolute deviation/std: standard deviation) of protein length and AFDB pLDDT as well as mean cluster size for representative structures (compare with Fig. 1B in the manuscript).

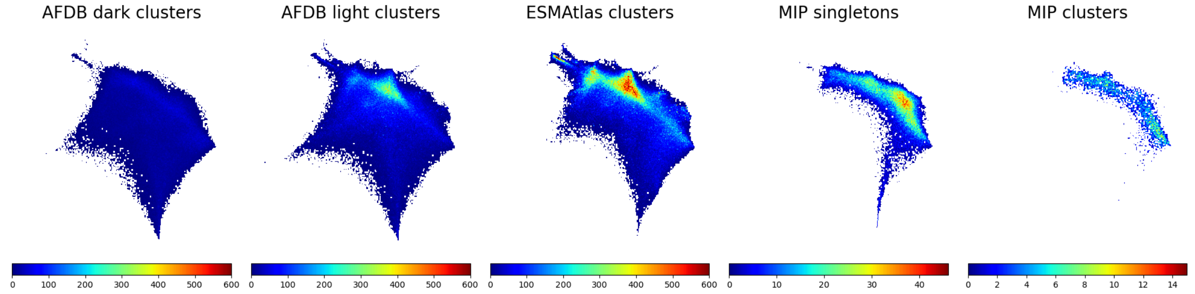

**Fig. S27:** The same as in Fig. 1C in the manuscript but the total number of structures (absolute values) is shown.

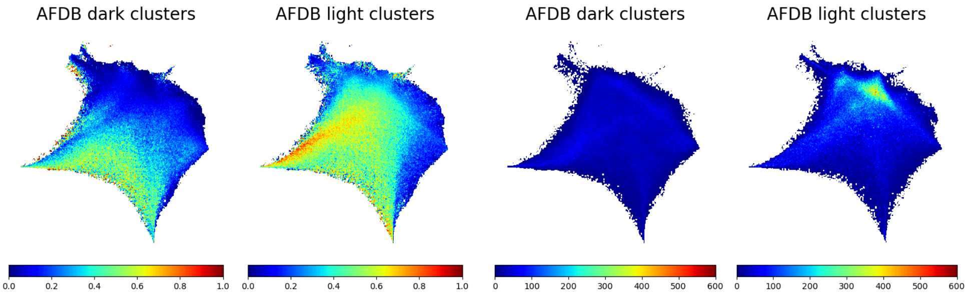

**Fig. S28:** The same as in Fig. 1C in the manuscript and Fig. S27 but for all AFDB models (two left plots: coverage, two right plots: total number of structures).

#### Plots for unnormalized Geometricus representations

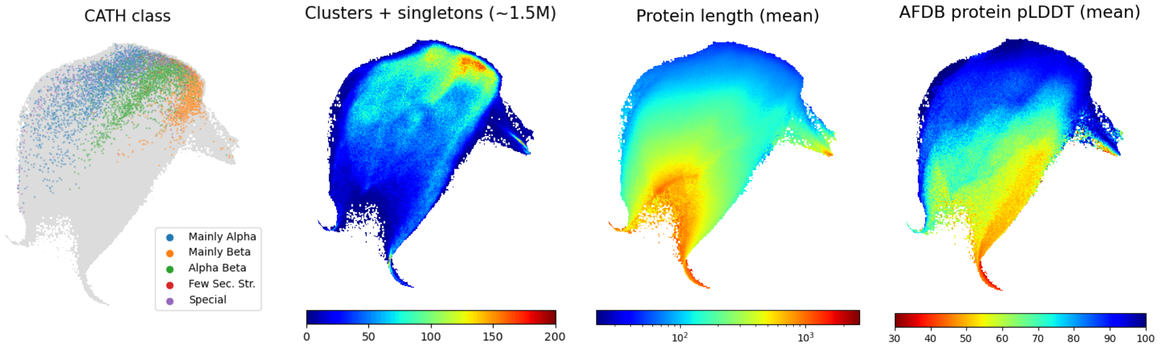

**Fig. S29:** The same as in Fig. 1B but for unnormalized Geometricus representations.

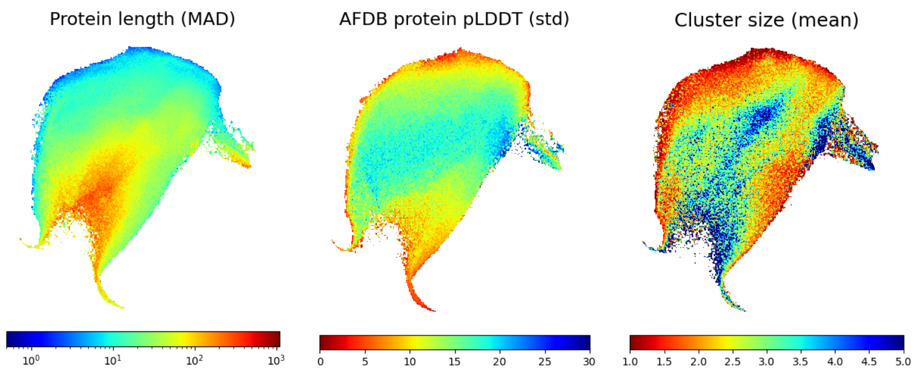

**Fig. S30:** The same as in Fig. S26 but for unnormalized Geometricus representations.

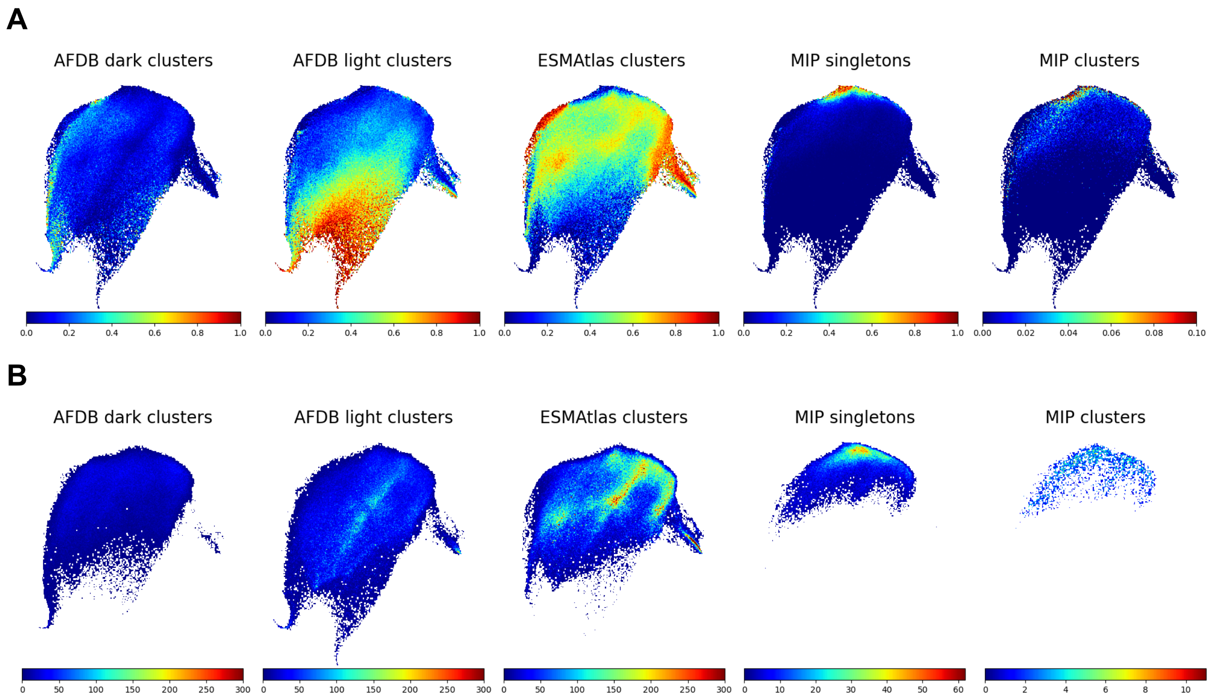

**Fig. S31: (A):** The same as in Fig. 1C in the manuscript but for unnormalized Geometricus representations. **(B):** Total number of structures (absolute values) is shown – compare with Fig. S27.

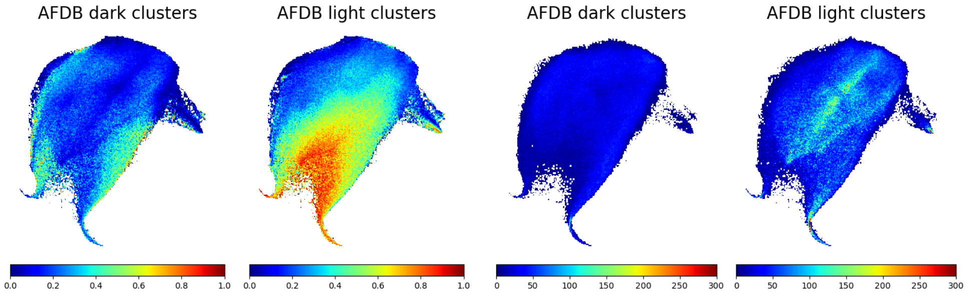

**Fig. S32:** The same as in Fig. S28 but for unnormalized Geometricus representations.

#### Other databases

To assess the generalizability of our dimensionality reduction approach, we evaluated it on two additional datasets: a smaller set of 10,000 AlphaFold2 models for sequences generated with ProtGPT2, and a larger set of 351,242 structural models from the Big Fantastic Viral Database (BFVD). For each structure in these datasets, we first computed Geometricus representations. Dimensionality reduction was then performed independently for each dataset (ProtGPT2 and BFVD) using PaCMAP, employing two distinct strategies:

1. Transformation using a pre-existing mapping – based on the AFDB, ESMAtlas, and MIP datasets – via the `transform()` method.
2. Fitting and transformation using an extended mapping – combining the AFDB, ESMAtlas, MIP, and the new dataset (ProtGPT2 or BFVD) – via the `fit_transform()` method.

The first approach is particularly useful when the goal is to project new structures into an existing embedding space – such as when identifying structural neighbors – though it's important to note that shapemers absent from the original Geometricus vocabulary are replaced with zeros. The second approach is more suitable when the objective is to understand how new datasets reshape or expand the existing structural landscape.

Fig. S33 and Fig. S34 illustrate the results of this analysis. For the ProtGPT2 dataset, proteins are distributed fairly evenly across the structural landscape, and minimal differences are observed between the two embedding strategies. In contrast, the BFVD dataset exhibits a more pronounced divergence between the two approaches. Using the transformation-only method (left panel of Fig. S34), the data reveals a concentration of alpha-helical proteins (visible in the lower corner). Viral proteomes comprise small single-helical proteins that, despite their structural simplicity, perform diverse critical functions during viral infection cycles. These compact elements contribute to viral assembly, membrane manipulation, immune antagonism, and host cellular pathway modulation. However, after fitting the embedding to include also the BFVD structures (right panel of Fig. S34) several notable changes become apparent:

1. Emergence of small, distant clusters – these outlier regions, labeled A–D, consist of short, low-complexity proteins.
2. Expansion of low-confidence region – the lower-left area, typically populated by low pLDDT structures, expands further due to the inclusion of additional low-complexity models (points E–H).
3. Displacement of alpha-helical cluster – the previously dense alpha-helical region is pushed outward from the main landscape (see area around point J).
4. Enrichment of a distinct region – a unique area at the top of the embedding space (surrounding point I) becomes enriched with BFVD proteins, predominantly long (>2000 amino acids) replicases and RNA-directed RNA polymerases.

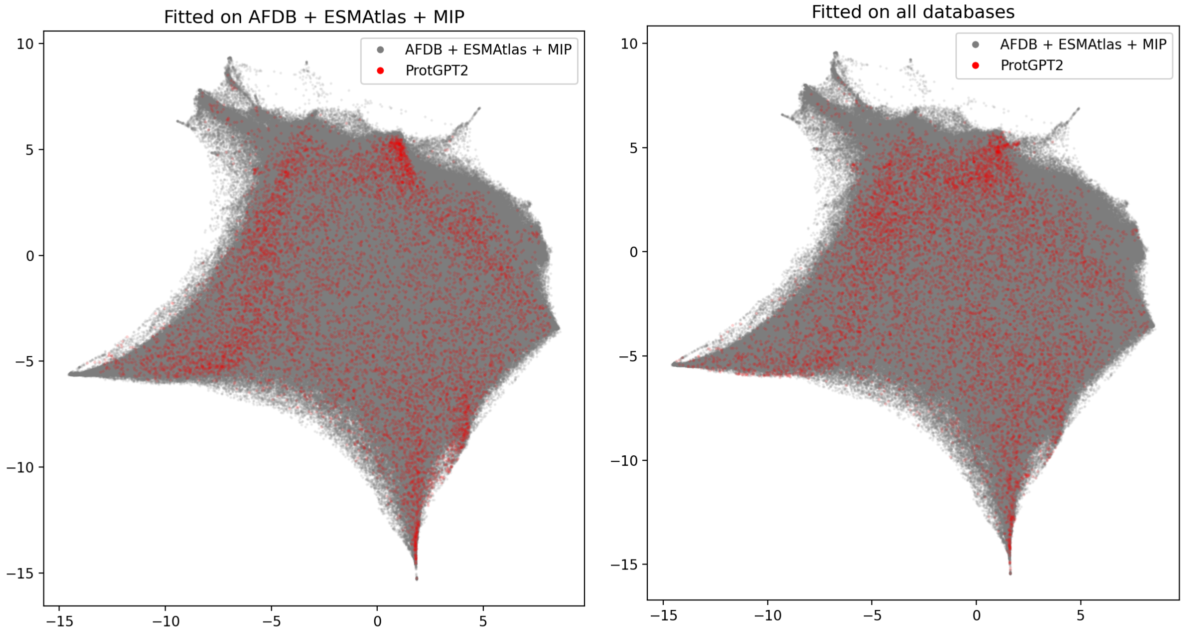

**Fig. S33:** 2-dimensional PaCMAP representation of 10,000 ProtGPT2-generated proteins, transformed using a PaCMAP model trained on AFDB, ESMAtlas, and MIP representative structures (left panel), as well as on all databases, including ProtGPT2 (right panel).

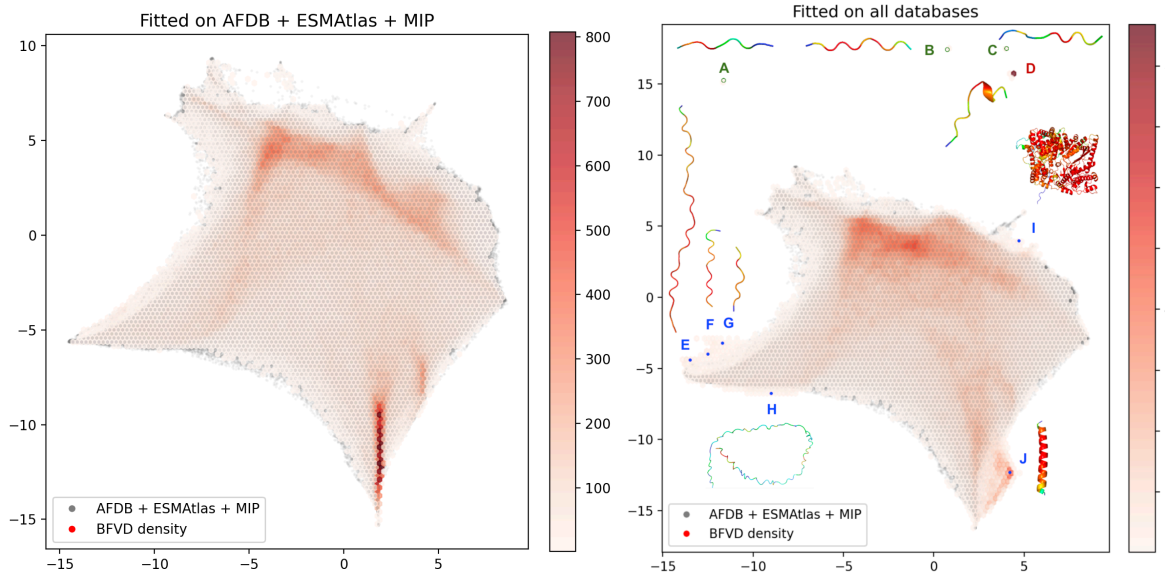

**Fig. S34:** 2-dimensional PaCMAP representation of 351,242 BFVD proteins, transformed using a PaCMAP model trained on AFDB, ESMAtlas, and MIP representative structures (left panel), as well as on all databases, including BFVD (right panel). Outlier points at the top of the right panel (labels: A, B, C) are highlighted with green circles for better visibility. Names of structural models in the right panel (labels: A-J) are shown in Table S3.

**Table S3:** Names of structural models presented in the right panel of Fig. S34.

| Point | Name |
|---|---|
| A | Q9IBY9_unrelaxed_rank_001_alphafold2_ptm_model_1_seed_000 |
| B | M1HRI5_unrelaxed_rank_001_alphafold2_ptm_model_1_seed_000 |

| | |
|---|---|
| C | F1ATE0_unrelaxed_rank_001_alphafold2_ptm_model_3_seed_000 |
| D | H9CID8_unrelaxed_rank_001_alphafold2_ptm_model_3_seed_000 |
| E | L7TJB1_unrelaxed_rank_001_alphafold2_ptm_model_2_seed_000 |
| F | M1PSG3_unrelaxed_rank_001_alphafold2_ptm_model_3_seed_000 |
| G | M9V8N1_unrelaxed_rank_001_alphafold2_ptm_model_3_seed_000 |
| H | A0A0F7L506_unrelaxed_rank_001_alphafold2_ptm_model_3_seed_000 |
| I | A0A2P1GNB0_1_unrelaxed_rank_001_alphafold2_ptm_model_3_seed_000 |
| J | A0A8S5P478_unrelaxed_rank_001_alphafold2_ptm_model_1_seed_000 |

To estimate how distinct the ProtGPT2 and BFVD datasets are from those used in our study, we applied foldseek easy-cluster to the original dataset of ~4 million proteins (prior to stage 2), supplemented with either 10,000 randomly generated structures from ProtGPT2 or 351,242 structures from BFVD. Clustering was performed using two parameter sets: (e-value = 0.01, coverage = 0.7) and (e-value = 0.001, coverage = 0.8). In both ProtGPT2 and BFVD, we observed a substantial number of singleton clusters, suggesting a high degree of structural uniqueness (see Table S4). For BFVD, this outcome was anticipated, given that viral proteins are scarcely represented in other databases (approximately 2% of the AFDB). To further explore the uniqueness of ProtGPT2-generated structures, we conducted foldseek easy-search against the AFDB, ESMAtlas, and MIP representative datasets using the `--greedybest-hits` option. The proportion of proteins with no detectable homologs was consistent with the clustering analysis: 73.3% at e-value = 0.001 and 69.6% at e-value = 0.01. To understand this lack of matches, we analyzed the mean pLDDT and pTM scores of the ProtGPT2 structures. Singletons exhibited significantly lower AlphaFold confidence scores, which may account for the absence of detectable homologs (see Fig. S35). Random selection of singleton and non-singleton ProtGPT2 models is presented in Fig. S37 (see also Table S5). The distribution of mean pLDDT scores in the BFVD dataset was more uniform. Notably, we identified a subset of high-confidence structures (mean pLDDT between 80 and 90), many of which correspond to alpha-helical proteins described above (see Fig. S36).

**Table S4:** Percentage of ProtGPT2 (first row) and BFVD (second row) singletons after the second stage clustering with foldseek easy-cluster. In both cases all 4,035,121 representative proteins from clustered AFDB, clustered ESMAtlas and MIP databases have been used (see Table S2).

| | **e-value = 0.01 coverage = 0.7** | **e-value = 0.001 coverage = 0.8** |
|---|---|---|
| **ProtGPT2** | 62.2% | 71.6% |
| **BFVD** | 53.9% | 64.1% |

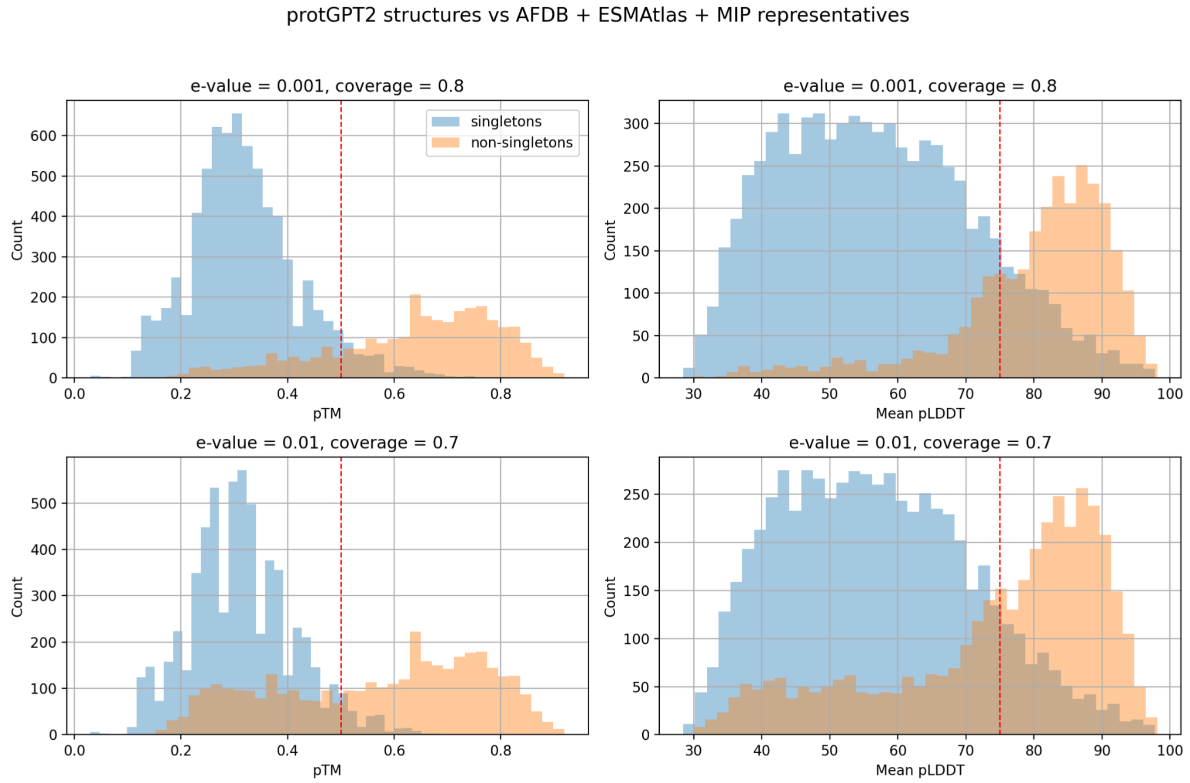

**Fig. S35:** pTM and mean pLDDT scores (left and right column respectively) for 10,000 ProtGPT2 protein structures without homologs (singletons after foldseek clustering) and with homologs (non-singletons). We used two sets of easy-cluster parameters (upper and lower panels respectively).

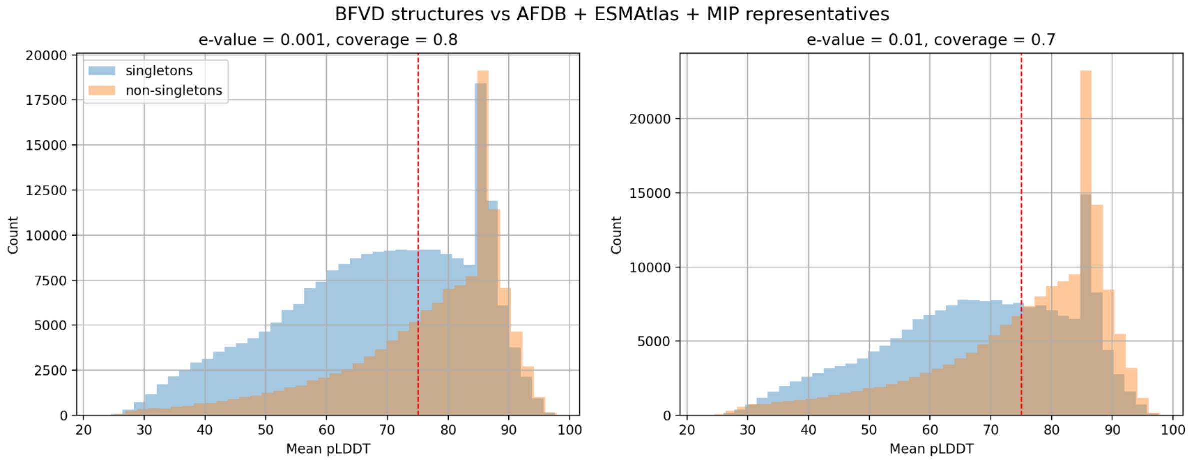

**Fig. S36:** Mean pLDDT for 351,242 protein structures without homologs (singletons after foldseek clustering) and with homologs (non-singletons). We used two sets of easy-cluster parameters (left and right panel respectively).

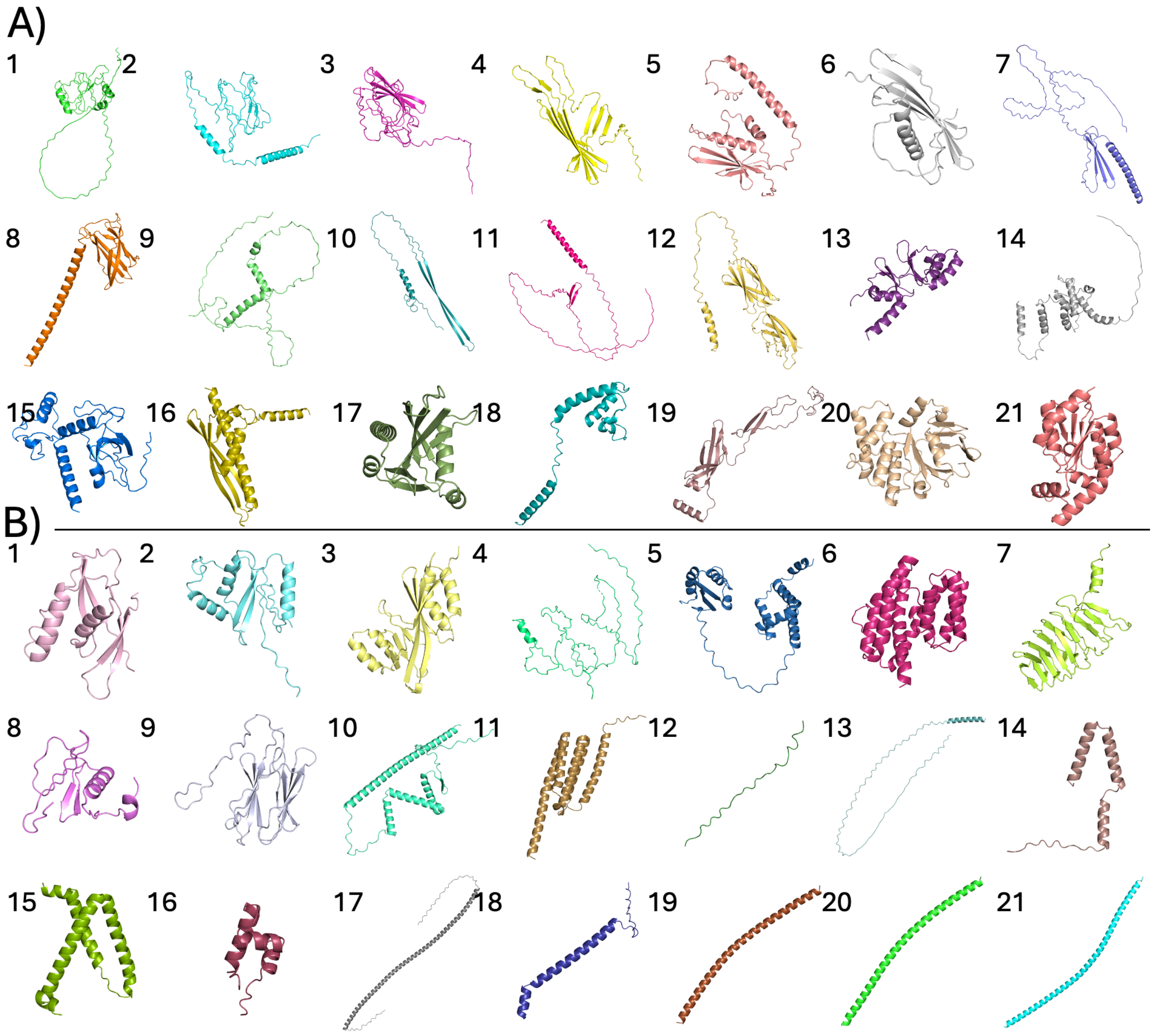

**Fig. S37**: Random selection of ProtGTP2 models (see Table S5), stratified by pLDDT range and cluster membership with 3 models per pLDDT scoring range. (A) Non-singletons and (B) singletons obtained from foldseek clustering with e-value = 0.001 and coverage = 0.8. Each group is subdivided into seven pLDDT intervals (30–40 to 90–100), enabling visual comparison of fold quality and diversity as a function of structural confidence and clusterability.

**Table S5:** Details of the models presented in Fig. S37.

| # | cluster | pLDDT range | model name |
|---|---|---|---|
| A1 | non-singleton | 30-40 | 3999_unrelaxed_rank_1_model_2.pdb |
| A2 | non-singleton | 30-40 | 6198_unrelaxed_rank_1_model_2.pdb |
| A3 | non-singleton | 30-40 | 6879_unrelaxed_rank_1_model_2.pdb |
| A4 | non-singleton | 40-50 | 3905_unrelaxed_rank_1_model_4.pdb |
| A5 | non-singleton | 40-50 | 5860_unrelaxed_rank_1_model_5.pdb |
| A6 | non-singleton | 40-50 | 6651_unrelaxed_rank_1_model_1.pdb |
| A7 | non-singleton | 50-60 | 4724_unrelaxed_rank_1_model_4.pdb |
| A8 | non-singleton | 50-60 | 7686_unrelaxed_rank_1_model_2.pdb |
| A9 | non-singleton | 50-60 | 8518_unrelaxed_rank_1_model_3.pdb |

| | | | |
|---|---|---|---|
| A10 | non-singleton | 60-70 | 2402_unrelaxed_rank_1_model_2.pdb |
| A11 | non-singleton | 60-70 | 2477_unrelaxed_rank_1_model_2.pdb |
| A12 | non-singleton | 60-70 | 3075_unrelaxed_rank_1_model_2.pdb |
| A13 | non-singleton | 70-80 | 2048_unrelaxed_rank_1_model_4.pdb |
| A14 | non-singleton | 70-80 | 2955_unrelaxed_rank_1_model_1.pdb |
| A15 | non-singleton | 70-80 | 3663_unrelaxed_rank_1_model_3.pdb |
| A16 | non-singleton | 80-90 | 1492_unrelaxed_rank_1_model_3.pdb |
| A17 | non-singleton | 80-90 | 1952_unrelaxed_rank_1_model_4.pdb |
| A18 | non-singleton | 80-90 | 586_unrelaxed_rank_1_model_3.pdb |
| A19 | non-singleton | 90-100 | 144_unrelaxed_rank_1_model_4.pdb |
| A20 | non-singleton | 90-100 | 206_unrelaxed_rank_1_model_5.pdb |
| A21 | non-singleton | 90-100 | 347_unrelaxed_rank_1_model_5.pdb |
| B1 | singleton | 30-40 | 6488_unrelaxed_rank_1_model_2.pdb |
| B2 | singleton | 30-40 | 7481_unrelaxed_rank_1_model_4.pdb |
| B3 | singleton | 30-40 | 9680_unrelaxed_rank_1_model_4.pdb |
| B4 | singleton | 40-50 | 3579_unrelaxed_rank_1_model_2.pdb |
| B5 | singleton | 40-50 | 4459_unrelaxed_rank_1_model_2.pdb |
| B6 | singleton | 40-50 | 6773_unrelaxed_rank_1_model_4.pdb |
| B7 | singleton | 50-60 | 4988_unrelaxed_rank_1_model_4.pdb |
| B8 | singleton | 50-60 | 7284_unrelaxed_rank_1_model_2.pdb |
| B9 | singleton | 50-60 | 8813_unrelaxed_rank_1_model_2.pdb |
| B10 | singleton | 60-70 | 3285_unrelaxed_rank_1_model_3.pdb |
| B11 | singleton | 60-70 | 4991_unrelaxed_rank_1_model_2.pdb |
| B12 | singleton | 60-70 | 9618_unrelaxed_rank_1_model_4.pdb |
| B13 | singleton | 70-80 | 3059_unrelaxed_rank_1_model_2.pdb |
| B14 | singleton | 70-80 | 6561_unrelaxed_rank_1_model_5.pdb |
| B15 | singleton | 70-80 | 9646_unrelaxed_rank_1_model_3.pdb |
| B16 | singleton | 80-90 | 198_unrelaxed_rank_1_model_3.pdb |
| B17 | singleton | 80-90 | 3674_unrelaxed_rank_1_model_3.pdb |
| B18 | singleton | 80-90 | 6062_unrelaxed_rank_1_model_5.pdb |
| B19 | singleton | 90-100 | 2837_unrelaxed_rank_1_model_3.pdb |
| B20 | singleton | 90-100 | 3281_unrelaxed_rank_1_model_5.pdb |
| B21 | singleton | 90-100 | 3669_unrelaxed_rank_1_model_3.pdb |

### Functional annotations

#### deepFRI v1.1

For training deepFRI v1.1 we utilized structures from the AlphaFold database v4 (https://alphafold.ebi.ac.uk/) and functional annotations from the Gene Ontology Annotation (GOA) Database (https://geneontology.org/). First, we selected GO terms with at least one experimental annotation and a total of 50 to 5000 annotations (excluding IEA). Additionally, we chose structures with ≥ 80% residues having pLDDT ≥ 70, and size between 60 and 1000 residues. This produced 2,822,622 structures annotated by 6,530 GO-terms and 4,905,062 annotations in total. Next, we enriched annotations by propagating GO-terms upwards in the GO-graph. This means that if a structure is annotated with a specific GO-term, it is also automatically annotated with all more general GO-terms linked to that term. This resulted in 8,877 GO-terms and 59,731,892 annotations in total. Lastly, the set has been randomly split into training and validation sets with the same proportions as in the original deepFRI release and three separate neural networks (for CC, MF and BP) have been trained with early stopping. Calibration curves showing precision versus deepFRI score (indicating how the method is certain about its predictions) for deepFRI v1.0 testing set are shown in Fig. S38. For both versions precision increases with the score but, at least for structure from the PDB, we should consider a higher deepFRI score for v1.1 as compared to v.1.0 in order to obtain similar quality predictions (e.g. 0.5 and 0.35 respectively for precision ≥ 0.5). We speculate that this may be caused by the fact that v1.1 has been trained with a much larger average number of structures per GO-term. Even though v1.1 produces more false positives for a given score, it predicts vastly more GO-terms as compared to v1.0 and may be especially useful for novel fold annotations (but manual curation is desired in such cases).

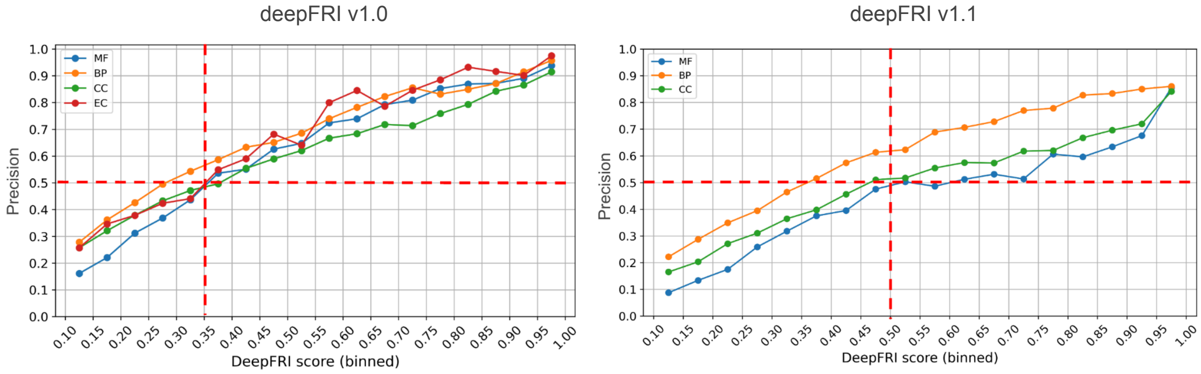

**Fig. S38:** Relation between precision and binned deepFRI score for releases 1.0 and 1.1 stratified by ontology (model), using deepFRI v1.0 testing set. The testing set comprises structures from the Protein Data Bank (PDB) with sequence similarity ≤ 30% to deepFRI v1.0 training set (see reference (4) for details).

#### Validation using E. coli proteome

We compared *E. coli* K12 proteome annotations derived from deepFRI v1.0, v1.1 with validated Swissprot records to benchmark our score thresholds – see Fig. S39. In general, deepFRI v1.0 is more accurate at scores ≥ 0.3, whereas deepFRI v1.1 shows higher concordance with Swissprot records at scores ≥ 0.5. However, annotations associated with superCOG 3 are less accurate in deepFRI v1.1 when compared to deepFRI v1.0.

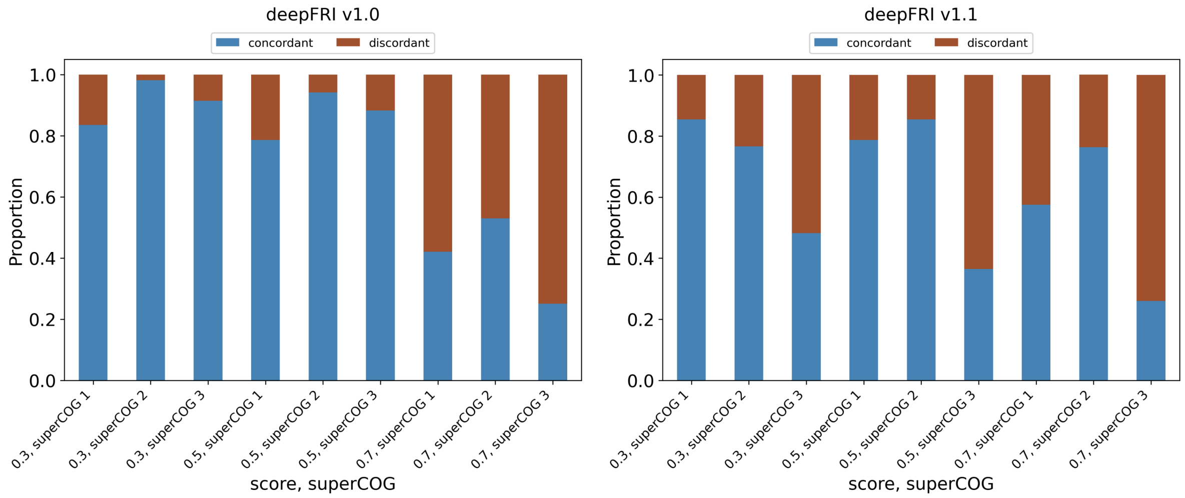

**Fig. S39:** Concordance and discordance plots (with respect to the Swissprot records) on superCOG level for *E. coli* K12 proteome. Benchmarks for deepFRI v 1.0 and v1.1 are shown for different score thresholds (0.3, 0.5 and 0.7).

Fig. S40 presents the aggregation of superCOG annotations for both deepFRI versions, Swissprot records, and the COG classifier ([https://github.com/moshi4/COGclassifier](https://github.com/moshi4/COGclassifier)).

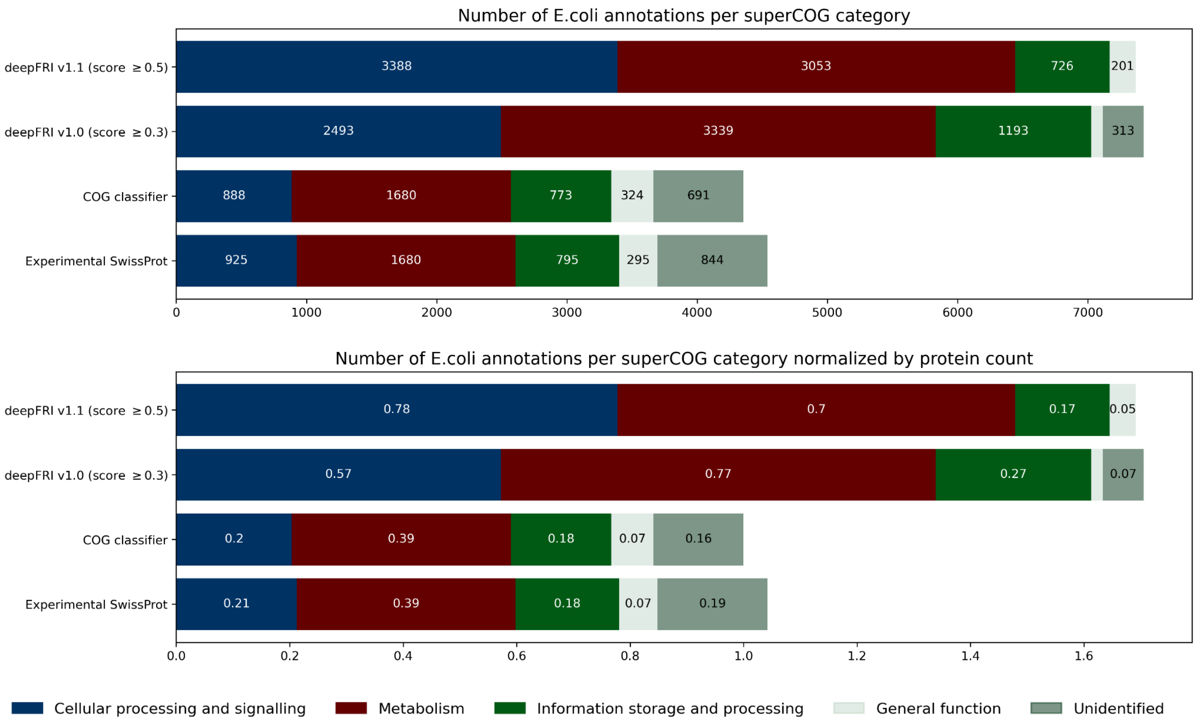

**Fig. S40:** Comparative analysis of *E. coli* K12 proteome annotations using deepFRI and COG classifier to Swissprot experimental records. Upper panel: total number of annotations. Lower panel: total number of annotations divided by the number of proteins. In the latter case, given the redundancy of annotations, more than one category can be assigned to a protein.

### Visualizations

Here, we present additional panels (supplementary to the main text) for visualizing 2D protein structure space using PaCMAP, for both normalized and unnormalized Geometricus representations, focusing on superCOG annotations.

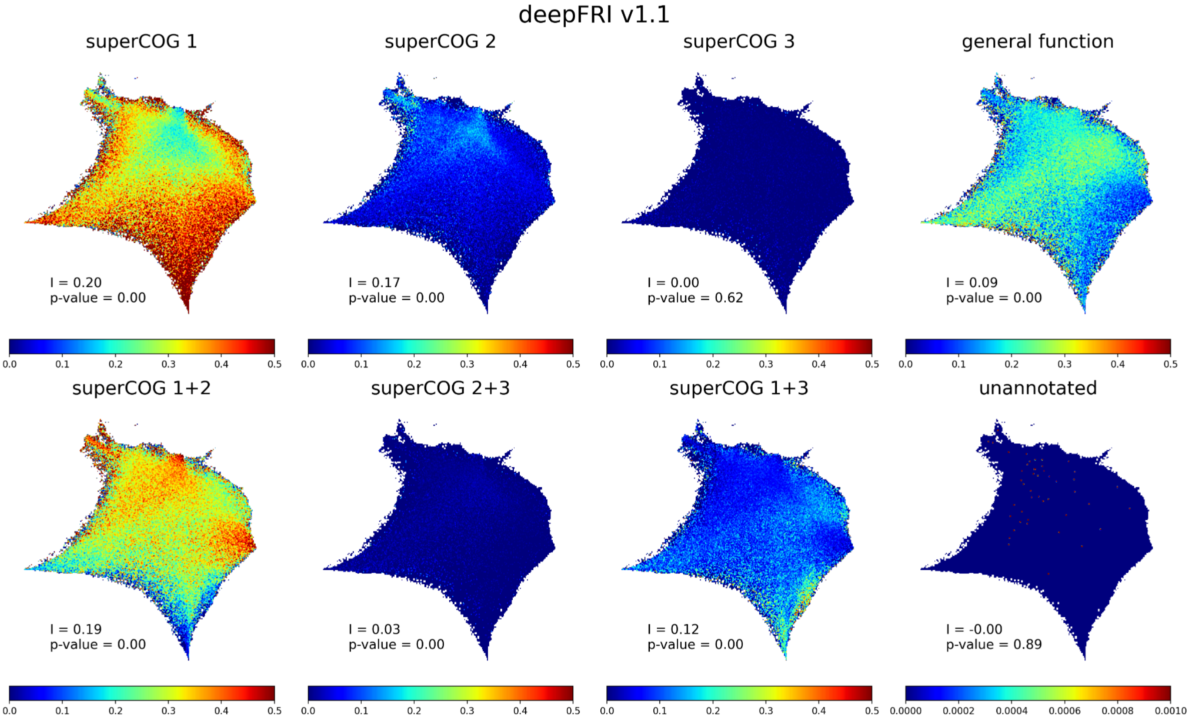

**Fig. S41:** The same as in Fig. 3B in the manuscript but for deepFRI v1.1.

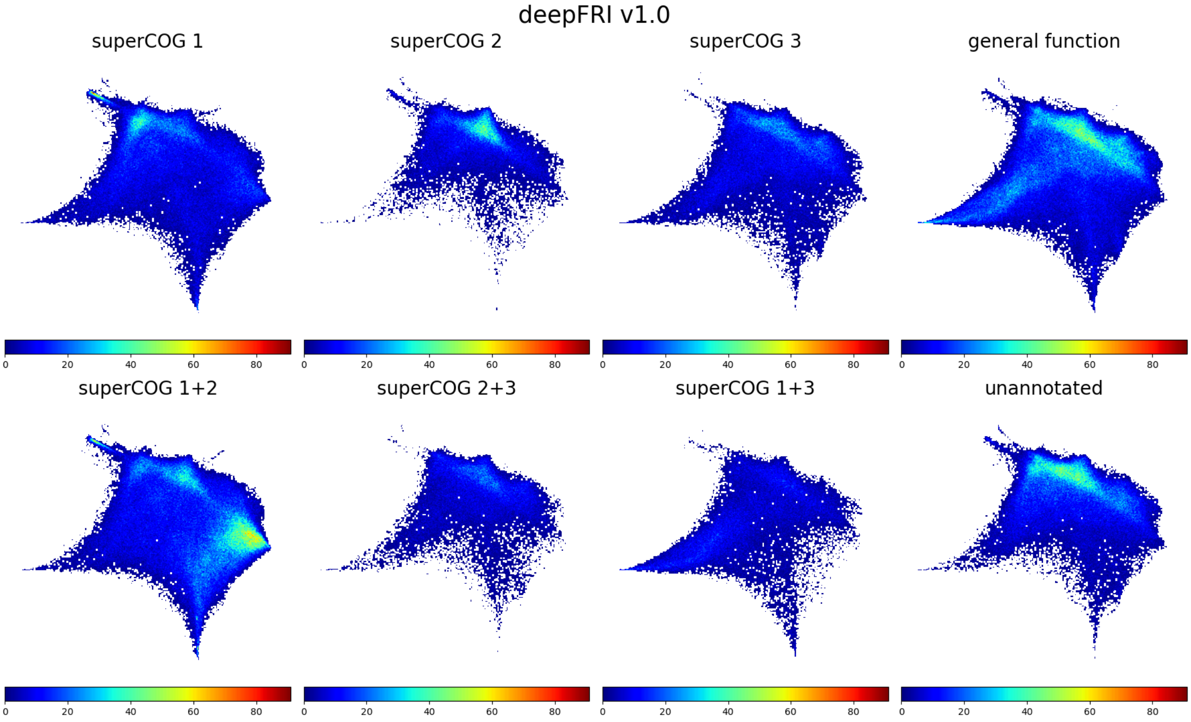

**Fig. S42:** The same as in Fig. 3B in the manuscript but the total number of structures (absolute values) is shown.

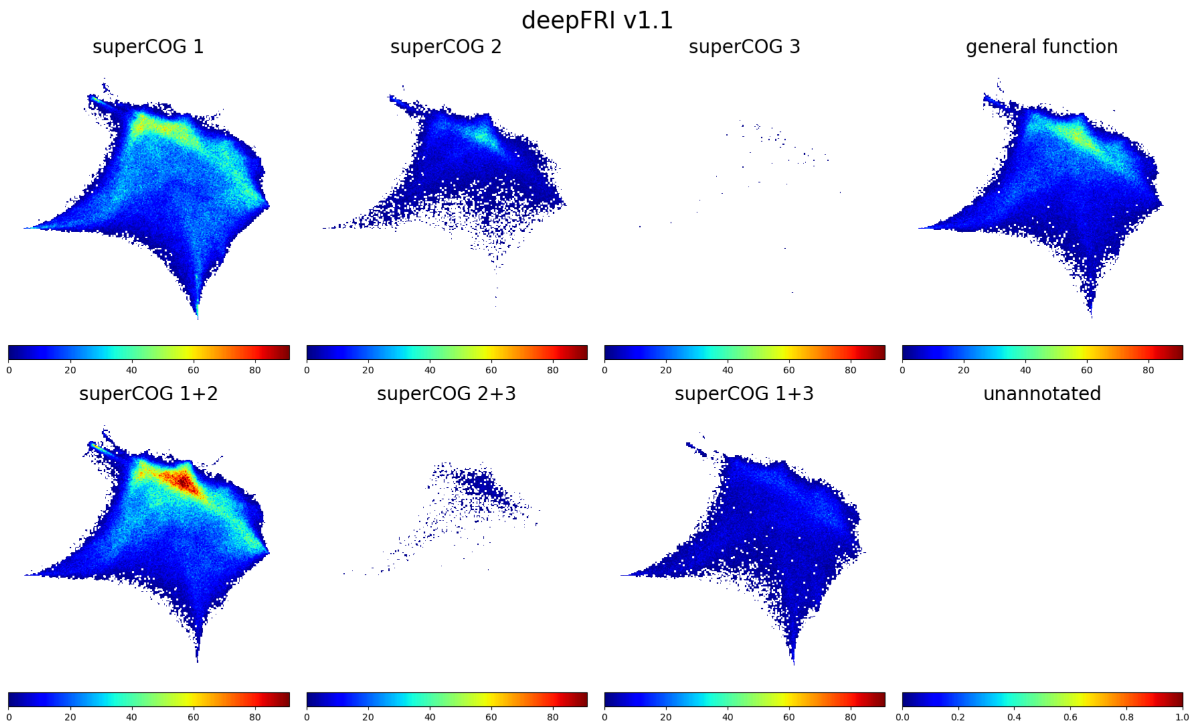

**Fig. S43:** The same as in Fig. 3B in the manuscript but the total number of structures (absolute values) is shown and deepFRI v1.1 is used.

#### Plots for unnormalized Geometricus representations

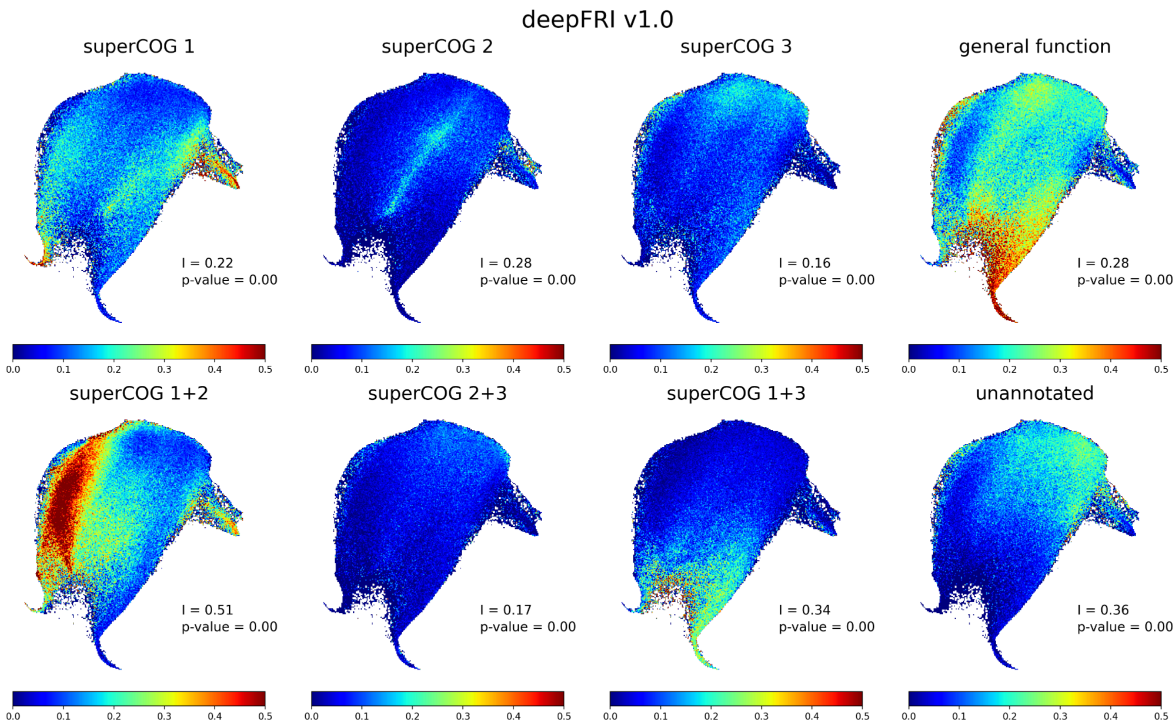

**Fig. S44:** The same as in Fig. 3B but for unnormalized Geometricus representations.

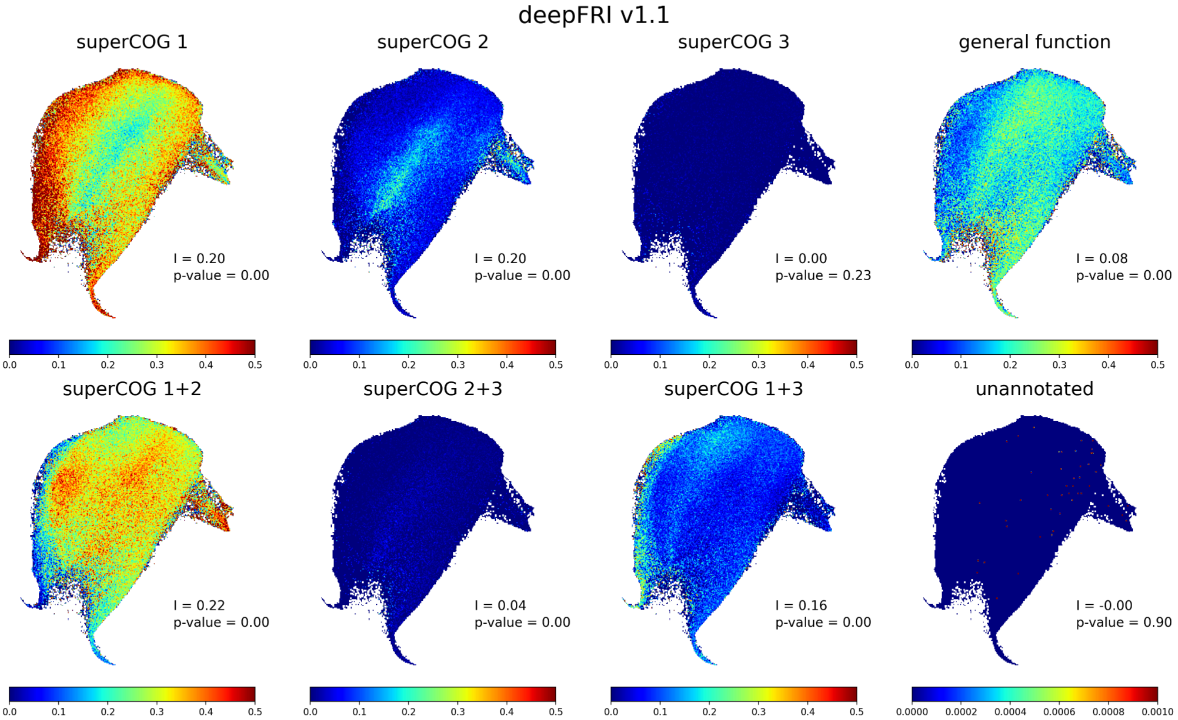

**Fig. S45:** The same as in Fig. Fig. S41 but for unnormalized Geometricus representations.

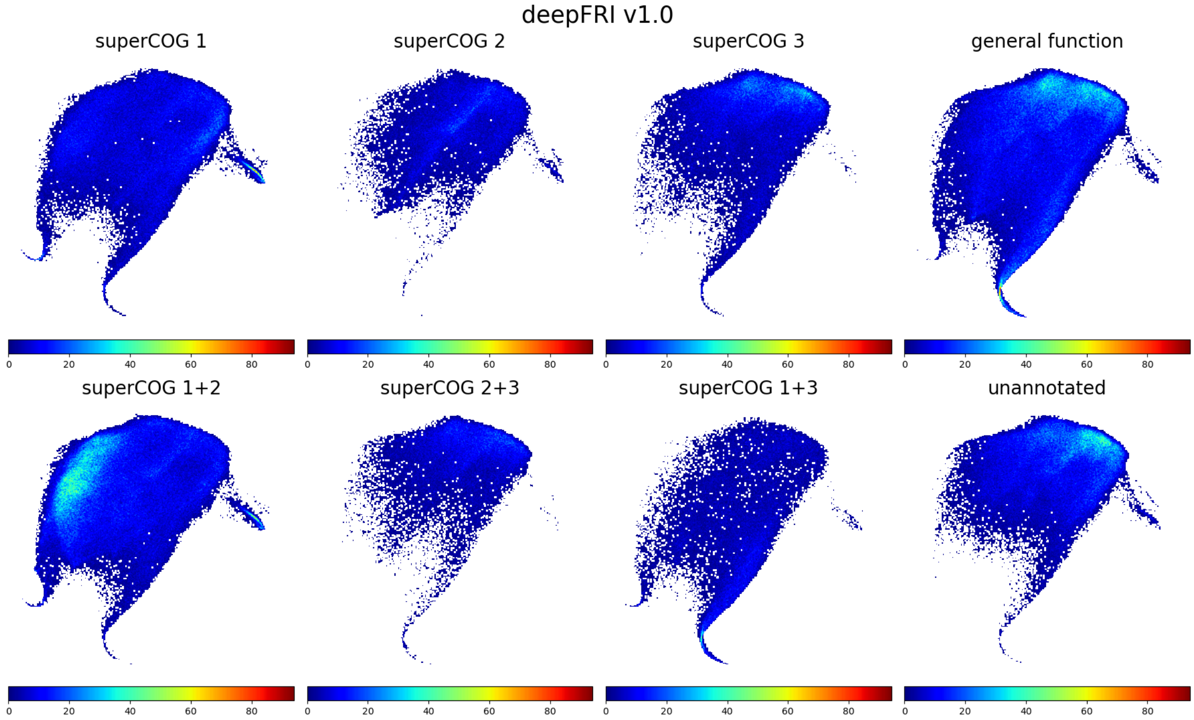

**Fig. S46:** The same as in Fig. Fig. S42 but for unnormalized Geometricus representations.

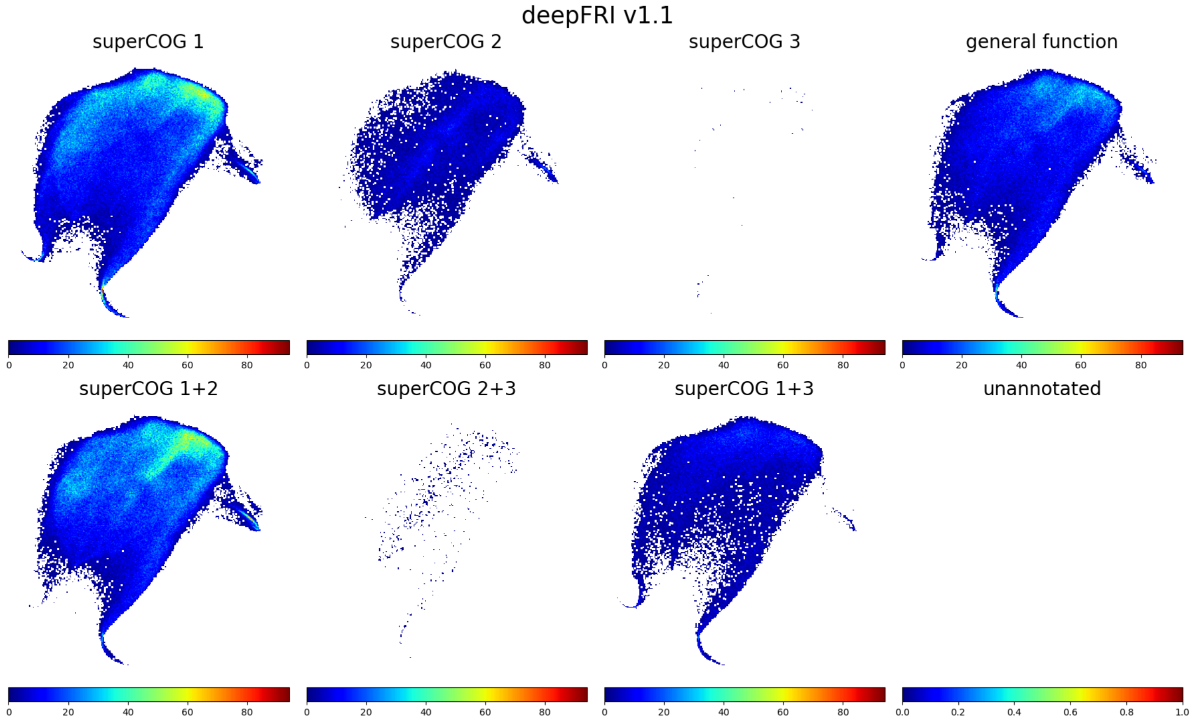

**Fig. S47:** The same as in Fig. Fig. S43 but for unnormalized Geometricus representations.

### Top COG categories

Overview of functional annotation profiles for each database, represented as COG categories abundance, can be seen in Fig. S48. Except for MIP new folds and AFDB dark clusters, the annotation profile shows very similar proportions within a given deepFRI version.

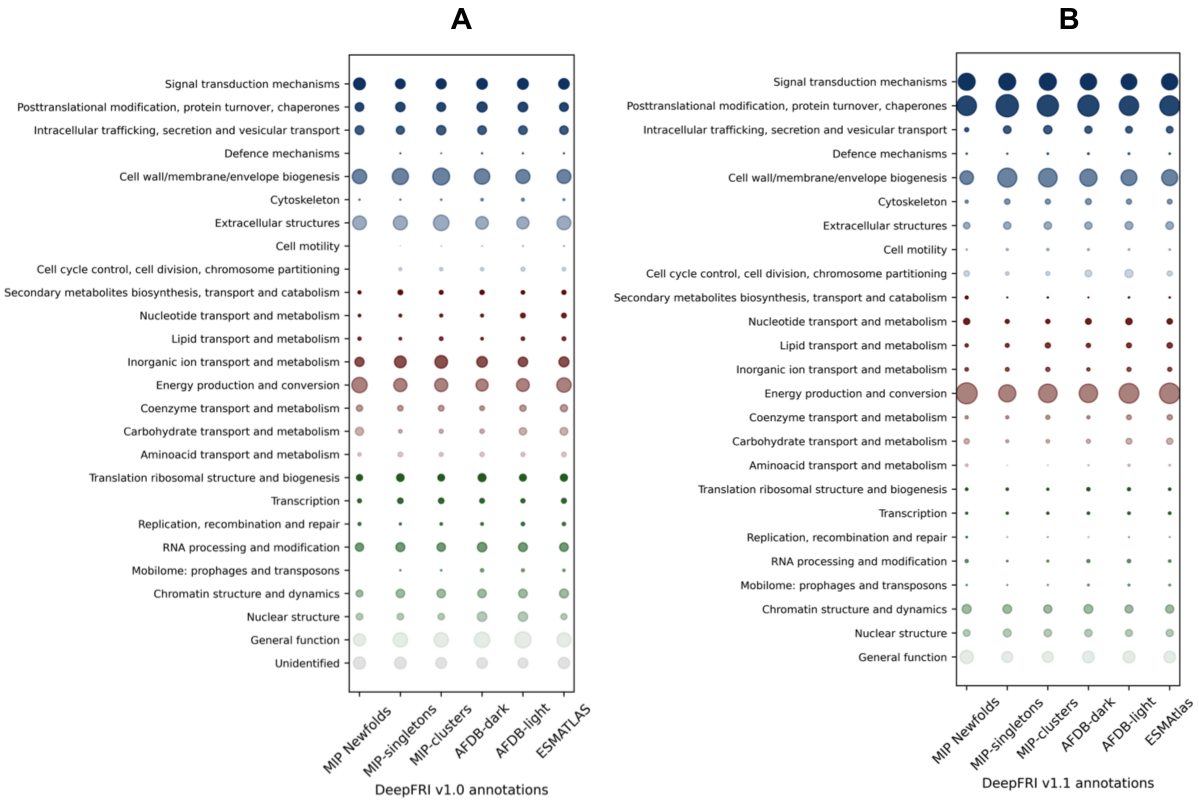

**Fig. S48:** Dot plot of proteins annotated by deepFRI v1.0 and v1.1 – **(A)** and **(B)** panels respectively. Annotations are represented as COG categories (with ≥ 1 COG per protein allowed) and dots represent relative abundance of COGs normalized to the protein count for each dataset.

### Cluster heterogeneity

In Fig. S49-Fig. S55 other variations of Fig. 5B-C are shown. Only high-quality AFDB models have been taken into account. In Fig. S49-Fig. S51, restricting the analysis to structures within a specific size range or using the same number of structures per cluster range, which are two potential sources of bias, does not alter the overall trends. Agreement between Fig. 5B and Fig. S49 is high for AFDB + ESMAtlas and slightly worse for AFDB + ESMAtlas + MIP (especially for superCOG 1) but at the same time the statistics is poorer (much less structures per a given cluster size range). The locations in the structure space of proteins forming heterogeneous clusters are shown in Fig. S56-Fig. S57, which correlates with regions of high overlap between AFDB light, ESMAtlas, and MIP structures (see Fig. 1C).

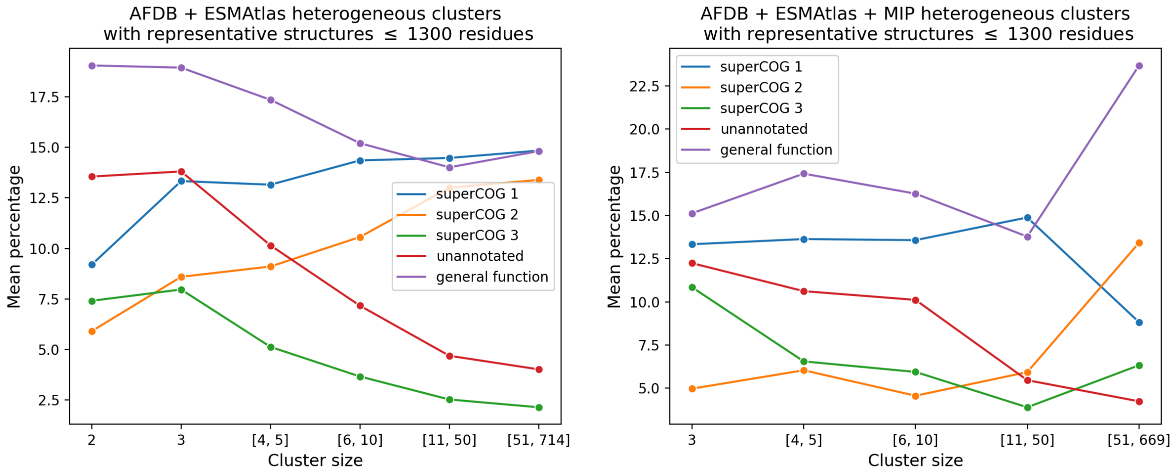

**Fig. S49:** Mean SuperCOG percentage for all structures in structurally heterogeneous clusters with high functional homogeneity (i.e., > 50% of structures annotated by one functional group). Percentage of clusters considered in the above plots: 35% and 28% for AFDB + ESMAtlas and AFDB + ESMAtlas + MIP respectively (compare with Fig. 5B in the manuscript where only representative structures are considered).

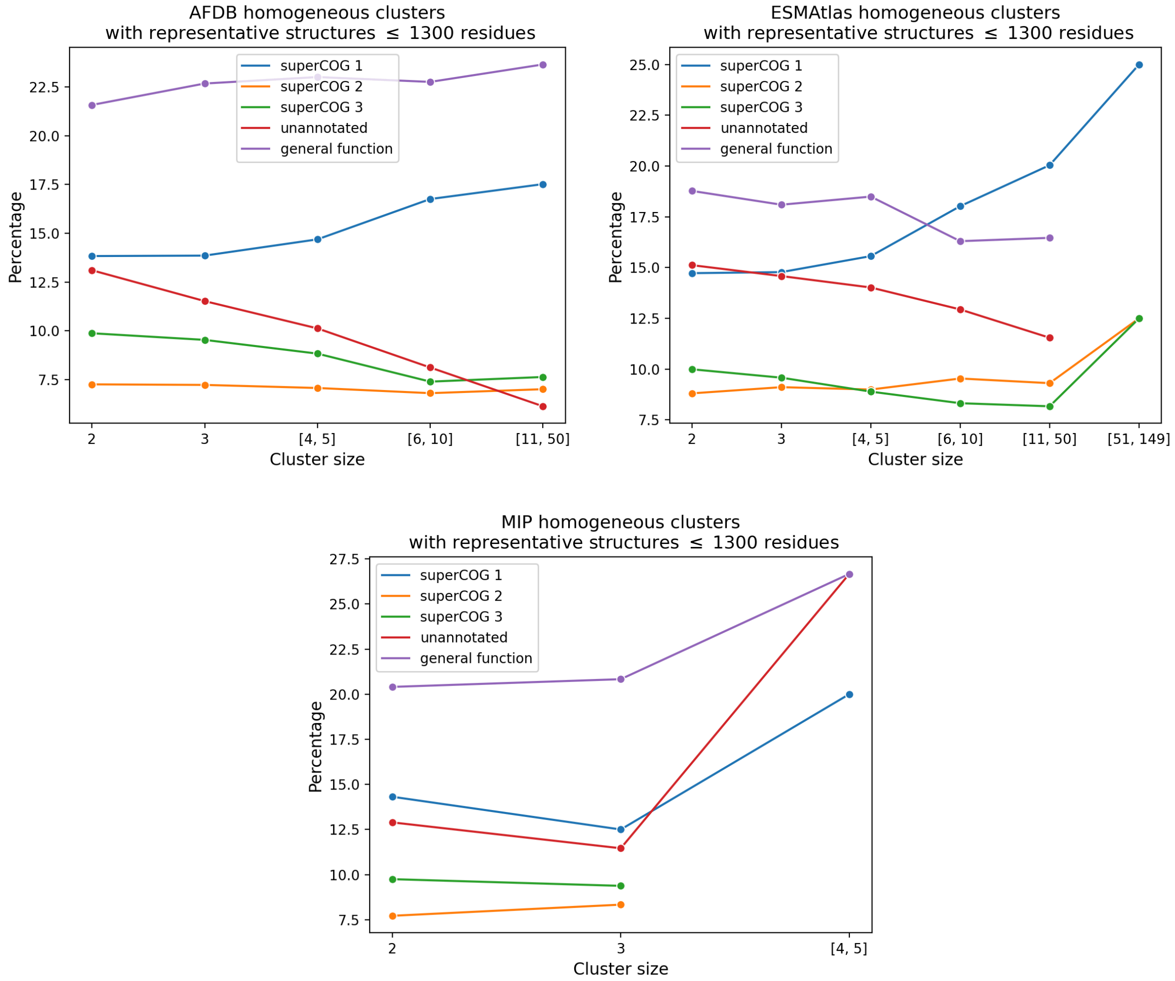

**Fig. S50:** The same as in Fig. 5B in the manuscript but for homogeneous clusters.

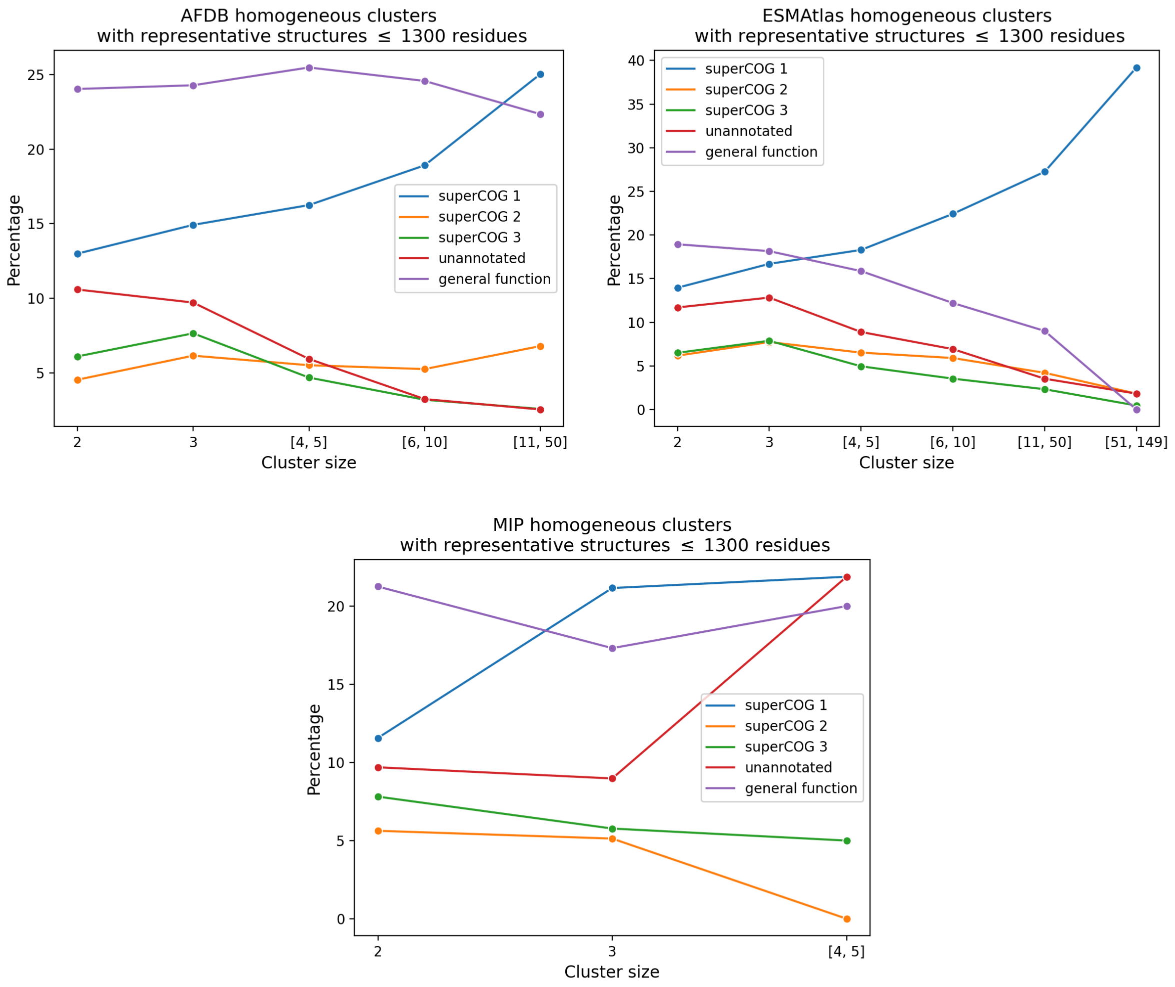

**Fig. S51:** The same as in Fig. S49 but for homogeneous clusters. Percentage of clusters considered in the above plots: 38%, 36%, 35% for AFDB, ESMAtlas and MIP respectively.

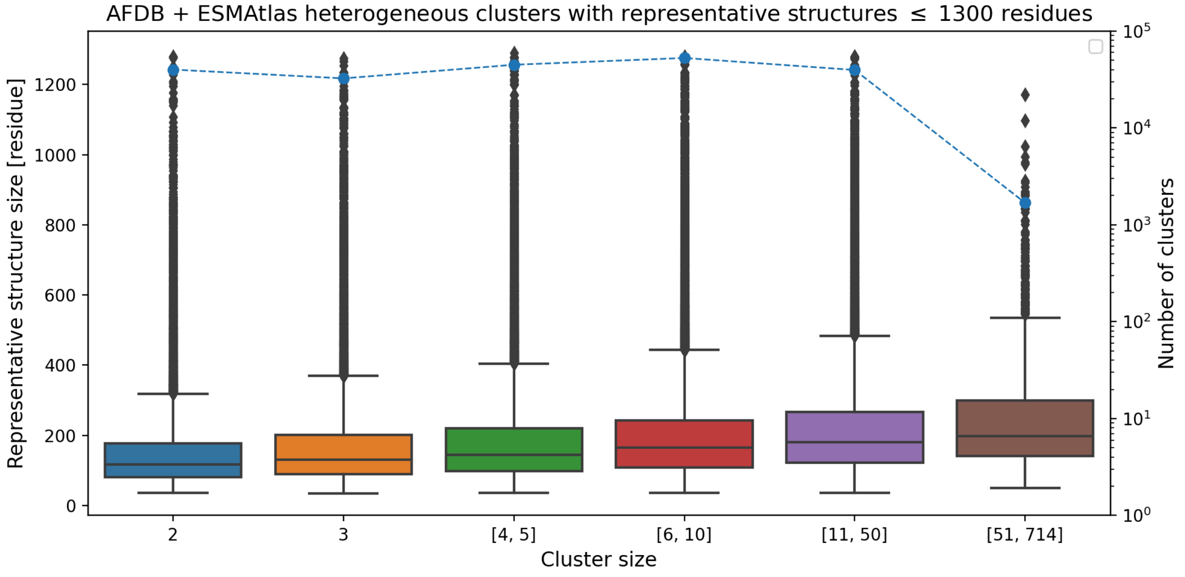

**Fig. S52:** The same as in Fig. 5C in the manuscript but for heterogeneous clusters having both AFDB and ESMAtlas structures.

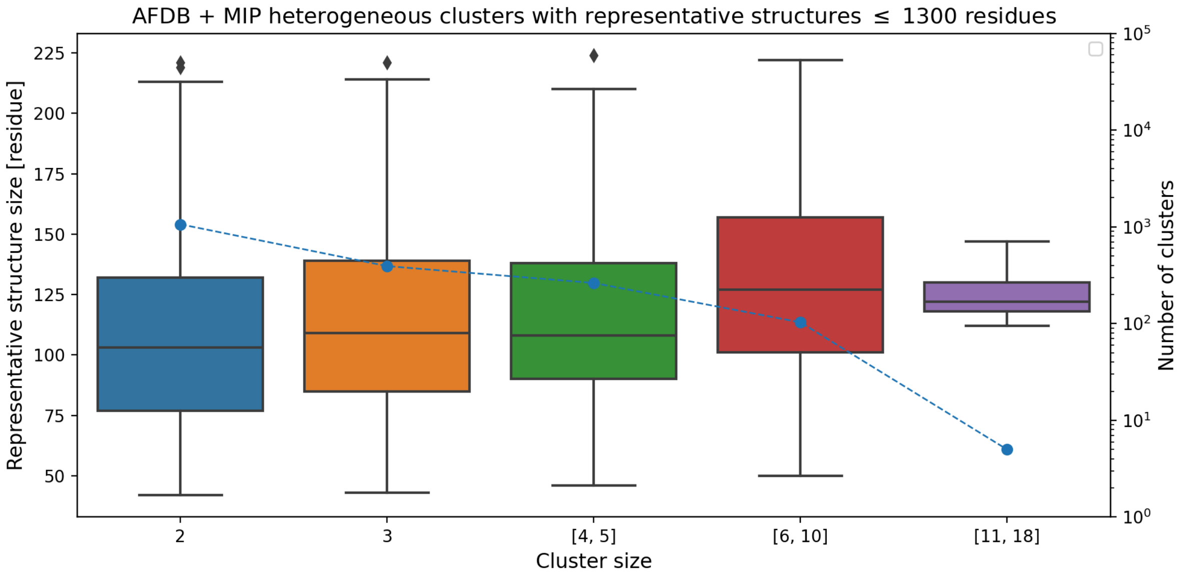

**Fig. S53:** The same as in Fig. 5C in the manuscript but for heterogeneous clusters having both AFDB and MIP structures.

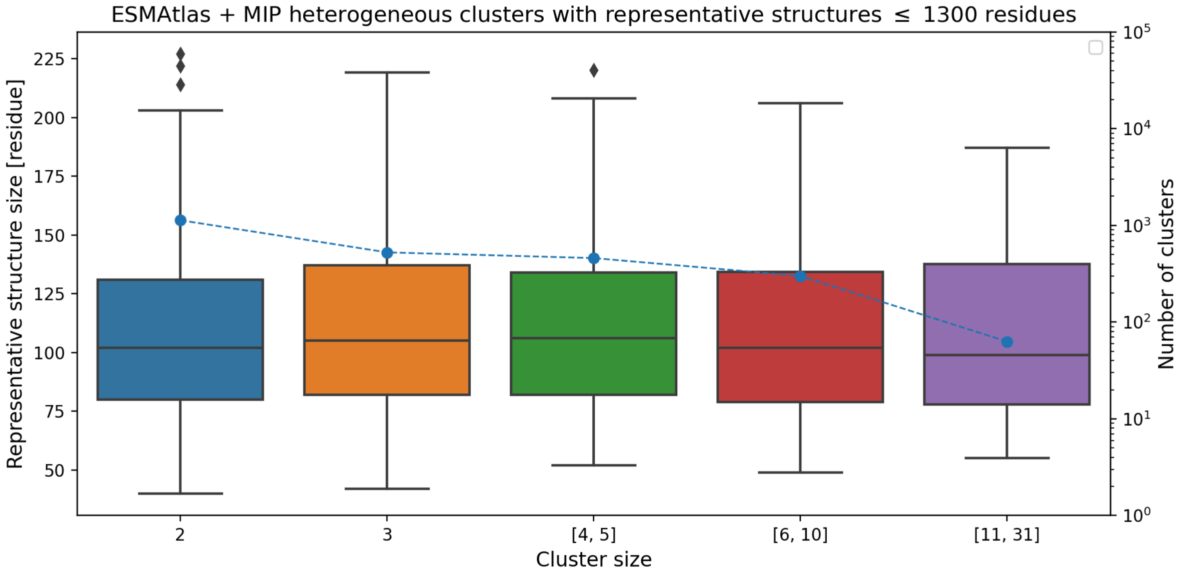

**Fig. S54:** The same as in Fig. 5C in the manuscript but for heterogeneous clusters having both ESMAtlas and MIP structures.

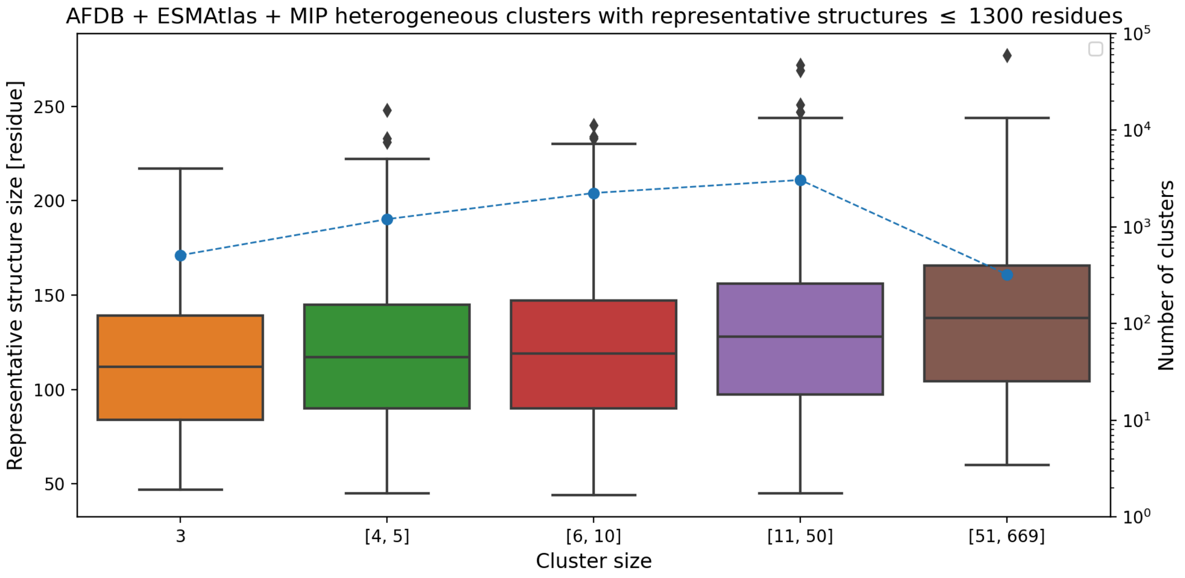

**Fig. S55:** The same as in Fig. 5C in the manuscript but for heterogeneous clusters having both AFDB, ESMAtlas, and MIP structures.

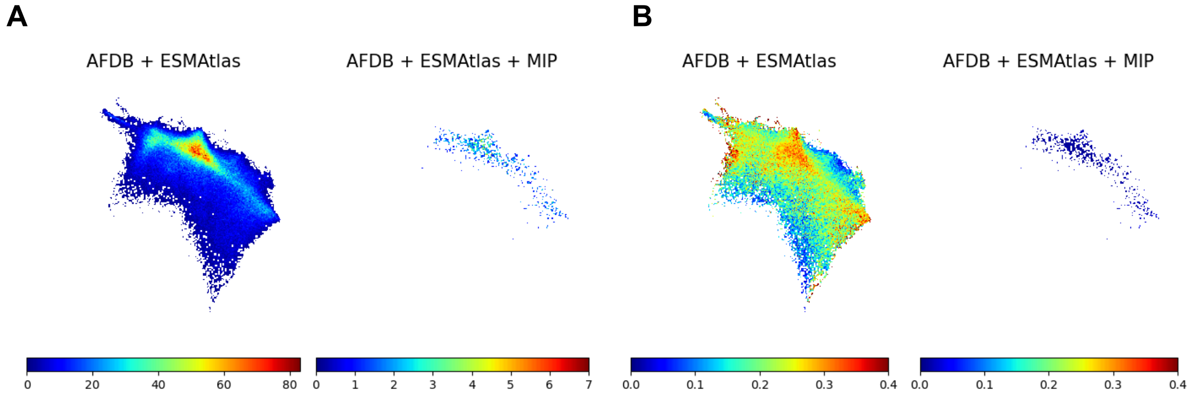

**Fig. S56: (A):** Number of representative structures originating from heterogeneous clusters of a given type (see titles). **(B):** Fraction of representative structures originating from heterogeneous clusters of a given type (see titles). Normalized Geometricus representations have been used. Only high-quality AF-DB models have been considered.

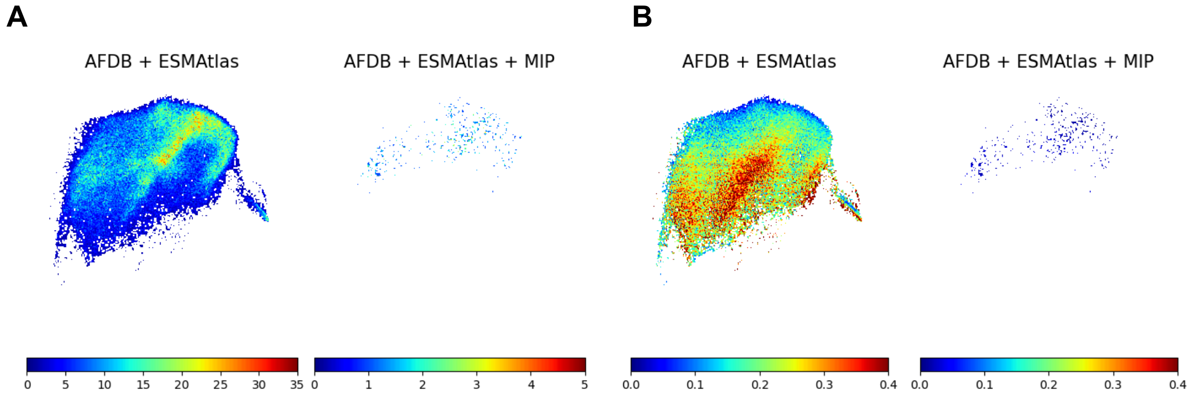

**Fig. S57:** The same as in Fig. S56 but for unnormalized Geometricus representations.

Fig. S58 highlights the largest representatives of homogeneous and heterogeneous clusters. Similarly to Fig. 2, we observe a high level of consistency across the structure space between different databases, although AFDB exhibits the greatest divergence among homogeneous groups. The heterogeneous clusters are located along the top edge of the space, with the AFDB + ESMAtlas group being more spread out and encompassing regions associated with ESMAtlas. Most of the structures in Fig. S58, represented in blue and magenta (groups A-F), resemble transmembrane or intramembrane proteins. This reflects both the ubiquity of these structures across databases and the diversity of cellular membranes among different organisms.

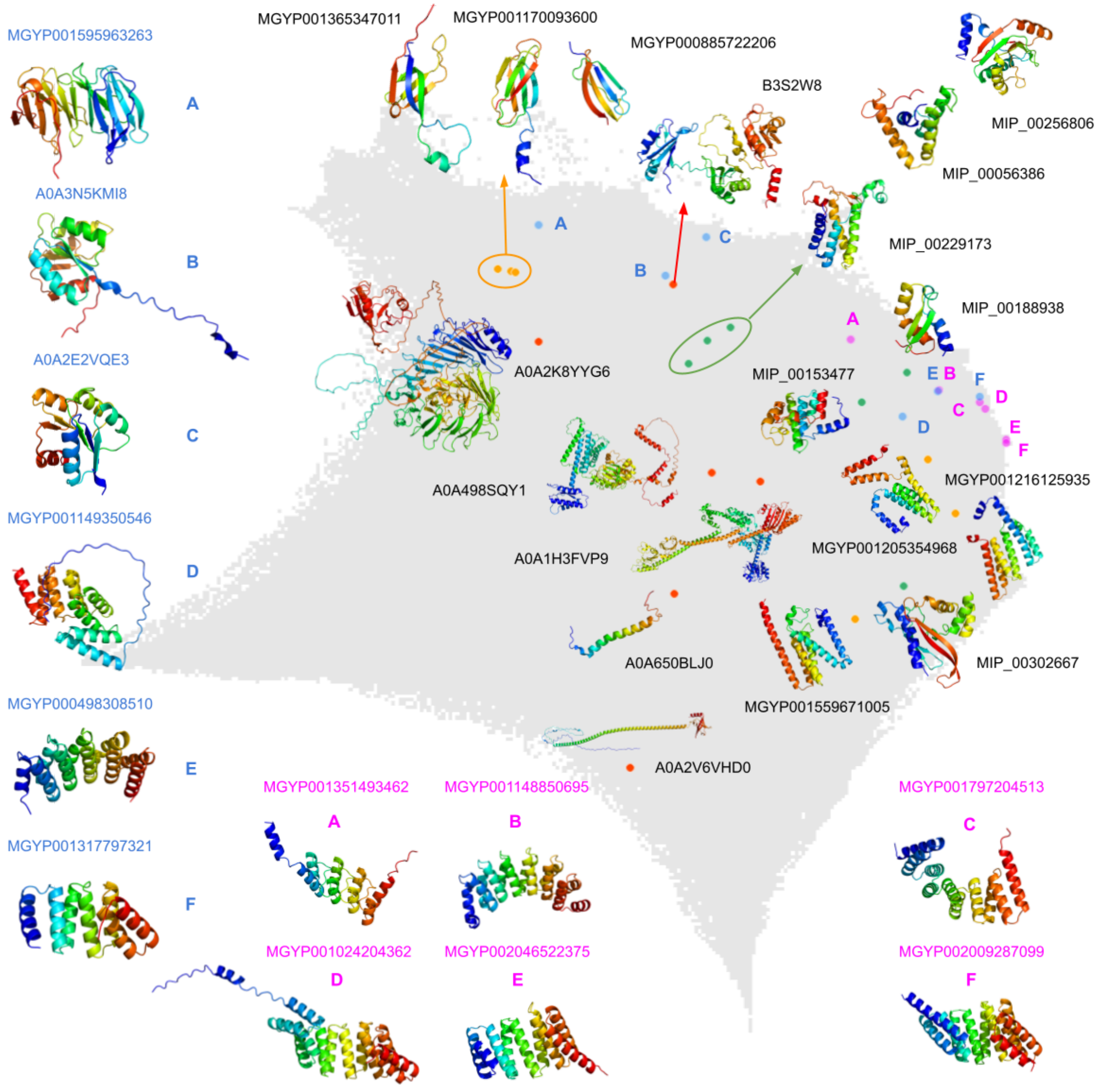

**Fig. S58: Largest cluster representatives.** Red, orange, and green dots represent AFDB, ESMAtlas, and MIP representatives of the top six largest homogeneous clusters, with names and structure visualizations attached to the dots on the plot. Similarly, blue and magenta dots correspond to representatives of the top six largest heterogeneous clusters for AFDB + ESMAtlas and AFDB + ESMAtlas + MIP, respectively, with names and structure visualizations displayed outside the plot. For easier navigation, blue and magenta dots are labeled with letters.

Fig. S59 provides a quantitative comparison of functions within homogeneous and heterogeneous clusters. The SuperCOG 1+2 category is the most prevalent (>23%) in AFDB + ESMAtlas homogeneous clusters and is also the most common in ESMAtlas homogeneous clusters (>23%). In contrast, the AFDB homogeneous cluster is dominated by the general function group (>22%) and SuperCOG 1+2 (>20%). For AFDB and ESMAtlas singletons, general function is the most prevalent, followed by SuperCOG 1+2, whereas this order is reversed in MIP singletons.

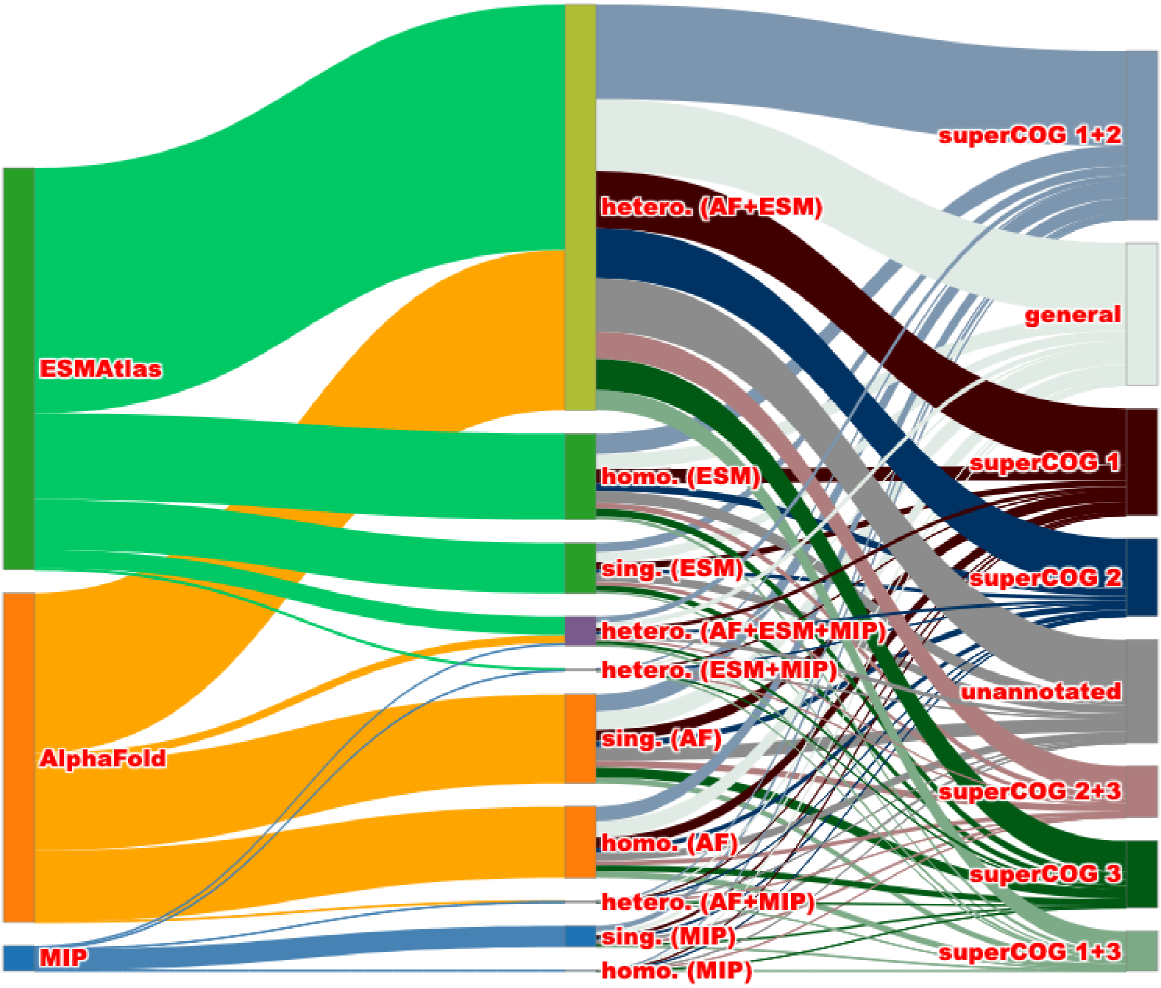

**Fig. S59:** Sankey diagram showing the number of AFDB, ESMAtlas, and MIP structures (first layer) forming different cluster types (second layer) and their functional annotation coverage (third layer).

### Taxonomy analysis

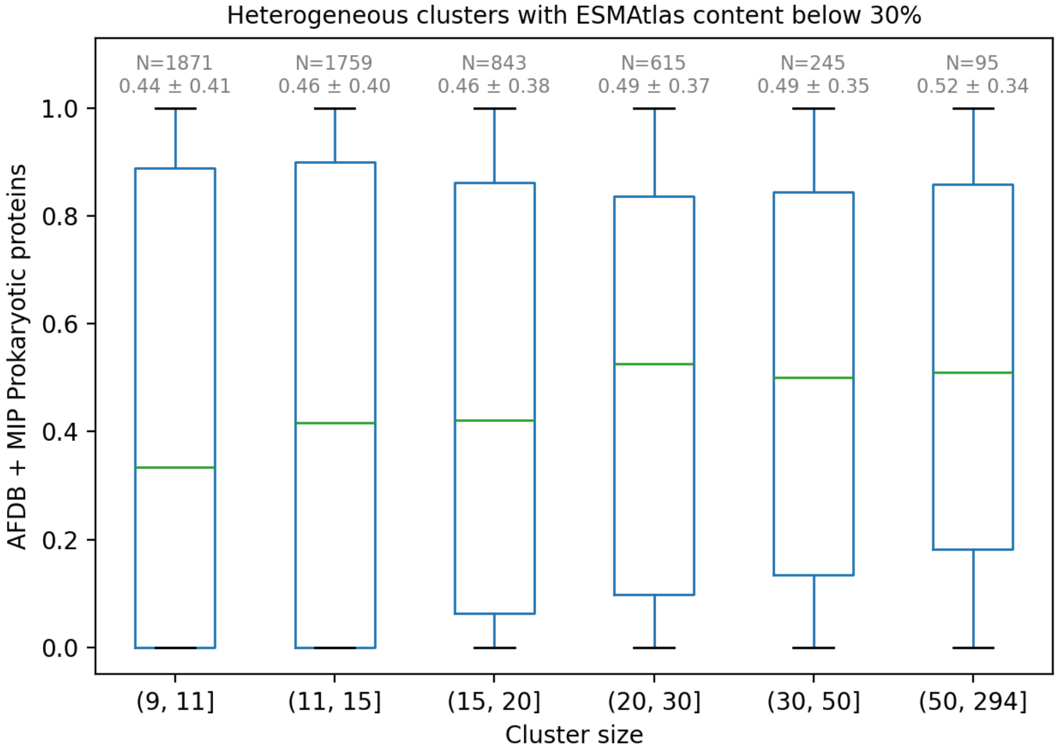

**Fig. S60:** Proportion of prokaryotic proteins with high quality AlphaFold models and MIP models in heterogeneous clusters with small ESMAtlas content (below 30%) as a function of cluster size. Above each box is the number of data points and mean +/- standard deviation.

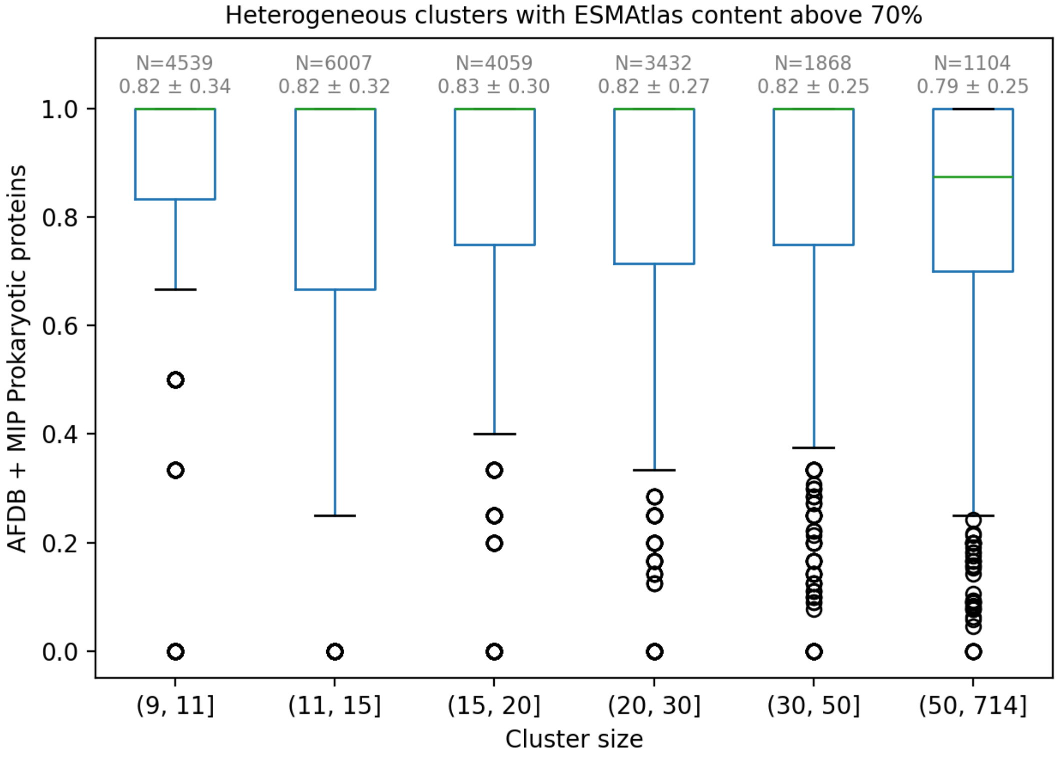

**Fig. S61:** Proportion of prokaryotic proteins with high quality AlphaFold models and MIP models in heterogeneous clusters with large ESMAtlas content (above 70%) as a function of cluster size. Above each box is the number of data points and mean +/- standard deviation.

**Table S6:** Number of proteins from each database for cluster representatives depicted in Fig. 6C.

| | **MGYP003387535279** | **MGYP001416922367** | **U6GJL0** | **A0A6A5UDC9** |
|---|---|---|---|---|
| AFDB light | 55 | 71 | 97 | 99 |
| AFDB dark | 0 | 1 | 0 | 0 |
| ESMAtlas | 78 | 71 | 48 | 348 |
| MIP | 0 | 0 | 0 | 0 |

**Table S7:** Taxonomic composition of cluster representatives used for entropy calculation (Fig. 6C).

| | **MGYP003387535279** | **MGYP001416922367** | **U6GJL0** | **A0A6A5UDC9** |
|---|---|---|---|---|
| Plants and Fungi | 20 | 30 | 34 | 48 |
| Bacteria | 5 | 12 | 20 | 24 |
| Invertebrates | 16 | 21 | 31 | 17 |
| Vertebrates | 9 | 4 | 7 | 5 |
| Primates | 0 | 2 | 2 | 2 |
| Environmental samples | 1 | 0 | 0 | 1 |
| Rodents | 1 | 0 | 1 | 1 |
| Mammals | 3 | 1 | 2 | 1 |
| Unknown | 0 | 2 | 0 | 0 |

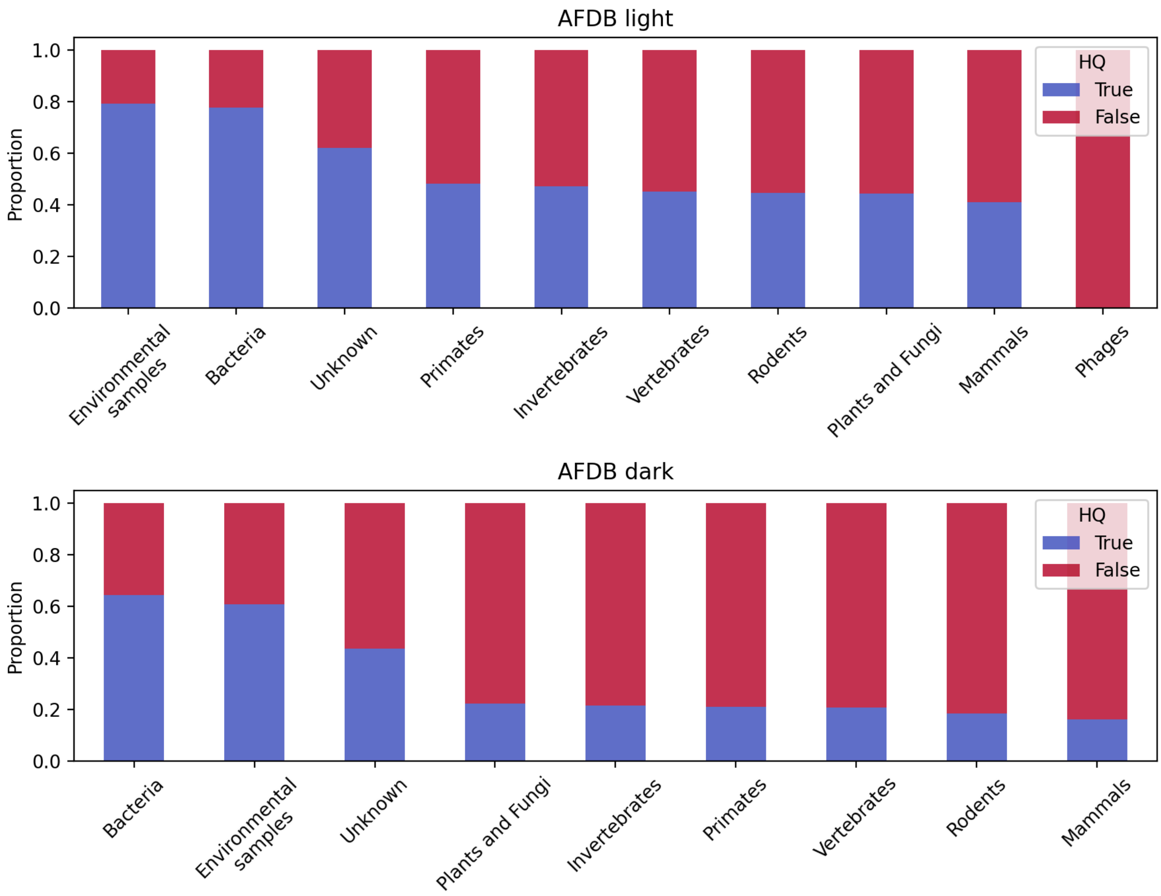

**Fig. S62:** Proportion of high quality AlphaFold predictions (HQ=True i.e. pLDDT > 70) stratified by taxonomic group for light and dark AFDB cluster representatives.
